# S-Phase induced RNAs control ORC1 engagement to H2A.Z and firing of early DNA replication origins

**DOI:** 10.1101/2021.10.19.465050

**Authors:** Simone Ummarino, Larysa Poluben, Alex K. Ebralidze, Ida Autiero, Yhanzou Zhang, Theodore Paniza, Madhura Deshpande, Johnathan Lee, Mahmoud Bassal, Bon Q. Trinh, Steven Balk, Robert Flaumenhaft, Jeannine Gerhardt, Sergei Mirkin, Daniel G. Tenen, Annalisa Di Ruscio

**Author notes:** These two authors equally contributed to the work (Larysa Poluben, Alex K. Ebralidze). These two authors equally contributed to the work (Daniel G. Tenen, Annalisa Di Ruscio). Contact: Annalisa Di Ruscio Center for Life Sciences 3 Blackfan Circle, Room 407, Boston, MA 02115; Daniel G. Tenen, Center for Life Sciences 3 Blackfan Circle, Room 437, Boston, MA 02115.

## Abstract

Coordinated initiation of DNA replication is essential to ensure efficient and timely DNA synthesis. Yet, the mechanism governing the “initiation” process in eukaryotic cells remains elusive. Here, we present data demonstrating a novel feature of RNAs transcribed in the proximity of actively replicating gene loci. We show that S-ph<u>A</u>se-RNAs a<u>NC</u>horing <u>OR</u>C1 (ANCOR*s*) to the histone variant H2A.Z are licensors of the DNA replication process. The concomitant ANCOR-H2A.Z interaction is essential for the cells to initiate duplication of their genetic heritage. Widespread and locus-specific perturbations of these transcripts correlate with anomalous replication patterns and loss of the replicative marker at the origin site.

Collectively, we unveil an RNA-mediated mechanism as the missing link for the generation of active replication origins and delineate a potential strategy to modulate replication in human cells at a local and global level.

## Introduction

In human cells, more than 6 billion base pairs of DNA need to be replicated and packaged into chromatin at every cell division cycle. Failures lead to mutations, epigenetic changes and chromosomal aberrations that ultimately cause genetic diseases and cancer [1].

While in prokaryotes the DNA replicates through well-defined single origin of replication, the DNA in eukaryotes forms multiple replication origins along the genome, the majority of which are not conserved among cell-types or tissues [2]. In yeast, autonomously replicating sequences (ARS), also known as “Replicators”, were shown to promote replication [3]. However, in mammals, only a handful of replication origins have been previously characterized [4–6], leaving the molecular mechanism controlling the distribution and activation of replication origins across diverse cell types and developmental stages relatively unexplored.

Human cell differentiation involves reproducible changes of DNA replication timing (RT), which is a cell-type specific feature, and refers to the temporal order whereby defined units of chromosomes replicate during the course of the S-phase in the cell cycle. Interestingly, the RT is disrupted in malignant transformation and parallel comparisons of RT profiles between normal and diseased tissues have enabled the identification of replication defects at specific gene loci: e.g. *c-MYC, FXN1* and *ROR1* that would not be identified by standard transcriptome analyses [7–9].

It was well established in studies of budding yeast and mammals, that the recognition of the origin site is mediated by a hexameric protein structure, termed the Origin Recognition Complex (ORCs)[10–13]). Recent developments in click chemistry-based approaches have facilitated the identification of multiple replication origins and confirmed the role of Origin Recognition Complex Subunit 1 (ORC1) in origin selection and replication timing [14]. Further, structural analyses by CryoEM have suggested a pre-licensing step wherein the interaction of ORC1 to a preassembled ORC2-5 complex promotes the opening of the ORC core and facilitates its DNA binding [15].

Yet despite multiple efforts, a sequence motif has not emerged from the analyses of ORC1-associated DNA, and interestingly, enrichment at enhancers and promoters marked by activating histone chromatin modifications has been reported for ORC1 [16, 17]. Together, these data have suggested that the plasticity of the mammalian DNA replication program could result from specific epigenetic features that modulates the selection and the activation of the replication origins. Recently, H. Long et al. have proposed a model wherein the histone variant H2A.Z recruits ORC1 at early replication origins in an H4K20me2-dependent manner [18]. However, this model fails to explain the determinants leading to enrichment of H2A.Z and coordinating ORC1 engagement at early S-Phase replicative origins. In this study we unveil the role of previously neglected participants, namely S-Ph<u>a</u>se RNAs a<u>NC</u>horing <u>OR</u>C1 (***ANCOR***) as licensor of mammalian replication origins, and describe a mechanism whereby *ANCORs* mediate the localization of ORC1 genome-wide while enabling binding of the histone variant H2A.Z, the major H2A variant associated with gene activation [19–21]. Importantly, perturbation of *ANCOR-H2A.Z* interaction results in changes of replication origin activity.

Our findings uncover an RNA-mediated mechanism linking genetic and epigenetic features of mammalian replication origins and define an uncharted mechanism to control replication timing in human cells.

## Results

### *c-MYC* S-Phase induced RNAs are implicated in c*-MYC* origin formation

Here, we investigated the contribution of S-phase induced RNAs in the initiation of replication origins. The rationale behind was twofold: 1) the coordination by cell-type-phase-specific RNAs would reconcile with the multiple replication origins observed across different cells and 2) the colocalization of H2A.Z and ORC1 at early replication origins would align with recent findings linking early S-Phase-induced RNAs to the deposition and modification of the histone variant H2A.Z [22].

First, we surveyed the *c-MYC* locus, the origin of which has been extensively studied [4, 6]. To examine the enrichment of ORC1 and H2A.Z at the *c-MYC* origin [6] hereafter referred as region 9-10; (**Figure 1a**), we carried out chromatin immunoprecipitation (ChIP) with the respective and validated antibodies, followed by quantitative (q)PCR (**Figure 1a**, **Supplementary Figure 1a**). We detected 4.7- and 14.5-fold enrichment within the region 9-10, for ORC1 and H2A.Z, respectively suggesting the presence of both proteins at the targeted *locus* (**Figure 1a**).

**Figure 1.**
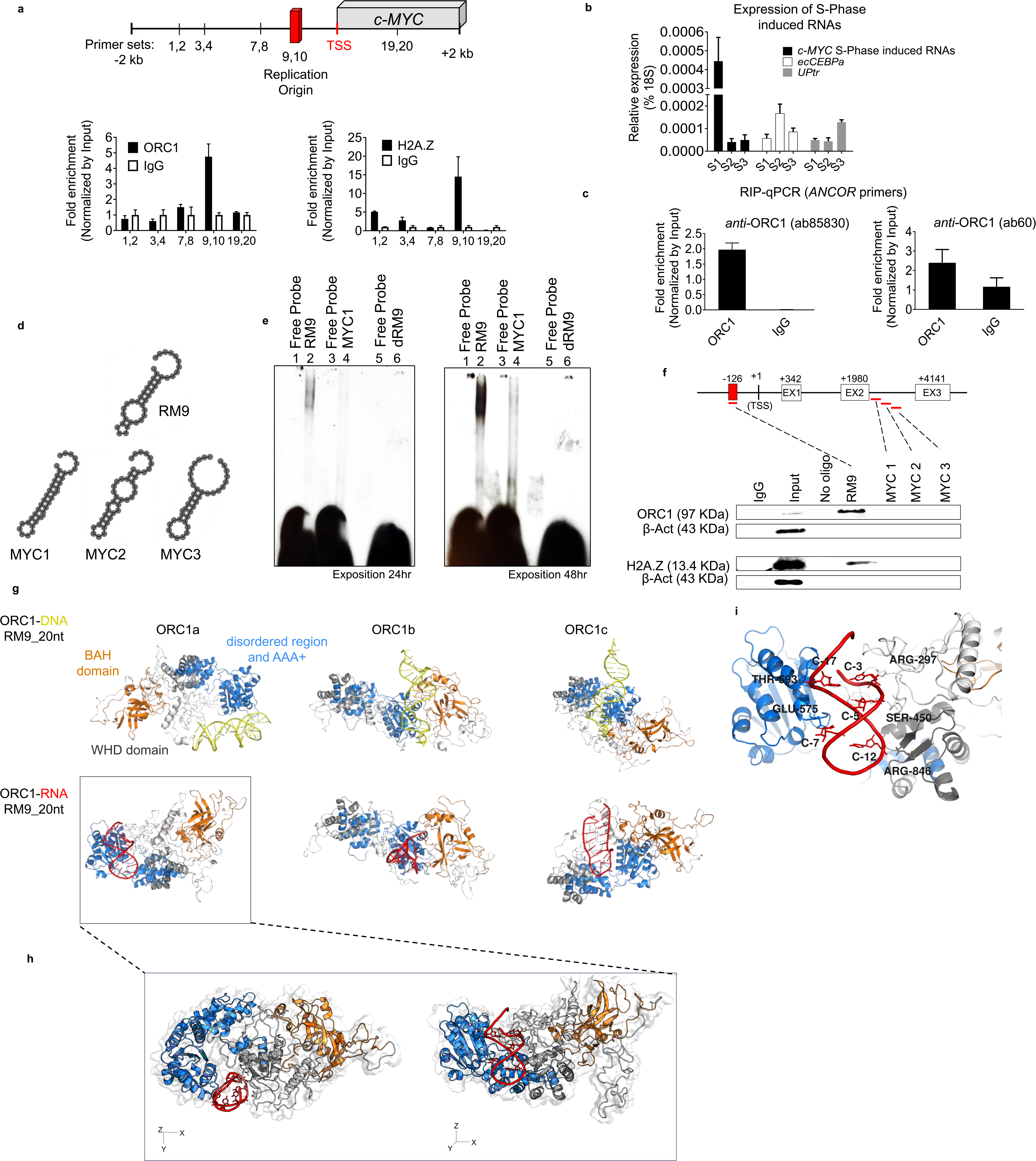
c*-MYC* S-Phase induced RNAs interact with ORC1. **a**. (**Upper panel**). Schematic of *c-MYC* locus and respective primer sets indicated by numbers. Red bar indicates *c-MYC* replication origin (region 9-10). (**Bottom panel**) ORC1 and H2A.Z enrichment at the *c- MYC* origin detected by chromatin immunoprecipitation (ChIP). The qPCR analysis was performed using distinct sets of primers designed as depicted in the schematic diagram provided above; **b**. Expression of S-Phase induced RNAs originated within the promoters of *c-MYC* and *CEBPA loci* at earlier and later stages of the S-Phase: S1(2h), S2(5h) and S3(8h), respectively; **c**. RNA immunoprecipitation (RIP) qRT-PCR performed using two separate antibodies against human ORC1 (ab85830 and ab60). The association between of *c-MYC* S-Ph**<u>a</u>**se RNAs a<u>nc</u>horing <u>OR</u>C1 – *<u>ANCOR</u>* is shown; **d-f**. *In vitro* RNA/DNA electrophoresis mobility shift assay (EMSA) and RNA pull-down assays. **d**. RNA secondary structures of the RNA probes corresponding to the *c-MYC* origin (RM9) and unrelated to the origin (MYC1, MYC2, and MYC3) calculated using RNAfold; **e**. RNA EMSA showing the preferential binding of ORC1 to RM9 (RNA) *versus* the unrelated MYC1 sequence and the homologous DNA sequence (dRM9); **f**. RNA pull down assay. Bound proteins were purified on streptavidin beads and immunoblotted with antibodies against ORC1 and H2A.Z. Experiment was repeated two times with similar results; **g**. Molecular dynamics simulations: frontal views representative structures of ORC1a, ORC1b and ORC1c states in complex with either dRM9 (DNA) or RM9 (RNA), respectively. RNA and DNA structures are derived using a RMSD based clustering approach and considering the last half of simulation time. All molecules are code-colored as follow, orange: ORC1-BAH domain (residues 1-200), marine ORC1 disordered region and AAA+ domain (residues 201-782), gray: orc1-WHD domain (residues 783-861); red and yellow: RM9 and dRM9, respectively; **h**. The figure shows a zoomed-in view of both the top and frontal perspectives, illustrating the preferred binding state of ORC1-RM9 complex; **i**. Zoom-in view showing the ORC1 ammino-acid residues interacting with RM9.

Second, we probed the nuclear fraction expression of S-Phase induced RNAs arising from the promoter and encompassing the *c-MYC* origin and others not arising from known replication origins during different stages of the S-Phase: S1(2h), S2(5h) and S3(8h) in K562 (Yildirim et al., 2020, Cell Reports 31, 107629) (**Supplementary Figure 1b**). A significant peak in the expression of the *c-MYC* S-Phase induced RNAs was detected in the early S-Phase in stark contrast with other RNAs not associated with origin of replications and showing higher levels at the later stages (S2 and S3) of the S-Phases (**Figure 1b**) [22, 23].

Importantly, the expression of the *c-MYC* S-Phase induced RNAs correlated with the expression profile of *c-MYC* mRNA (**Figure 1b**, **Supplementary Figure 1c**).

The precise mechanisms guiding the human (h)ORC1 localization on chromatin and its replication function remain poorly understood [24]. It was shown that RNAse A treatment impairs ORC1 recruitment *in vitro* to the Epstein–Barr virus plasmid replication origin [24] and similarly the interaction between telomeric repeat-binding factors 2 (TRF2) and ORC1 [25]. Recent studies further indicated an ORC1 preferential binding for G-rich RNA or single-stranded DNA sequences *in vitro* [26] overall suggesting a potential contribution of RNA in the ORC1 recruitment. Therefore, we explored the possibility of an RNA-dependent mechanism coordinating the initiation and the chromatin status at replication origins in eucaryotic cells. To determine whether ORC1 could physically associate with the *c-MYC* S-Phase induced RNAs, we performed RNA immunoprecipitation (RIP) with two different ORC1 antibodies (**Figure 1c**).

We observed enrichment of *c-MYC* S-Phase induced RNAs in the ORC1-RNA precipitates and termed these transcripts *c-MYC ANCOR* (S-Ph**<u>A</u>**se RNAs a**<u>NC</u>**horing **<u>OR</u>**C1). To study the molecular properties of RNA-ORC1 interaction *in vitro*, we performed RNA/DNA electrophoresis mobility shift assay (EMSA) and RNA pull down assay with K562 cell extracts (**Figures 1d-f**, **Supplementary Figure 1d**). The RNA oligonucleotides were selected based upon the: (1) proximity to the *c-MYC* origin and the higher CG content (RM9); and (2) distance from the *c-MYC* origin and the lower CG content (MYC1,2,3). DNA oligonucleotides (dRM9) corresponding to the RM9 sequence were also analyzed to compare the RNA and DNA interaction with ORC1. RNA-ORC1 complex formation was observed with RM9 (**Figure 1e**, left and middle panels of **Supplementary Figure 1d**). By contrast, no binding with the homologous DNA oligonucleotide was detected suggesting a non-preferential interaction of ORC1 for DNA as compared to RNA (**Figure 1e**, left and middle panels of **Supplementary Figure 1d**). RNA oligonucleotides unrelated to *c-MYC* origin failed to produce a shift by RNA EMSA (**Figure 1e**, left and middle panels of **Supplementary Figure 1d**) and did not produce ORC1 enrichment by RNA pull down assay (**Figure 1f**), suggesting the binding specificity between RM9-ORC1. Strikingly, we confirmed the presence of H2A.Z in the RM9 precipitates by RNA pull-down suggesting to coexistence of both ORC1 and H2A.Z in the complex (**Figure 1f**). Consistently, the interaction between H2A.Z and RM9 but not MYC1 was also detected by EMSA (right panel **Supplementary Figure 1d**), supporting the dual role of RNA in coordinating localization of ORC1 while enabling binding to the histone variant H2A.Z [22].

Next, we sought to analyze *in silico* the interaction between the (h)ORC1 and the *c-MYC ANCOR* with respect to the equivalent DNA sequence in order to understand the structural and dynamic features of ORC1-RNA complexing. The interaction between ORC1 and RNA (RM9) and the respective DNA sequence (dRM9) was modeled using theoretical methods. Three apo ORC1 states (ORC1a, ORC1b and ORC1c) were considered as equally probable conformations targeted by the sequence (**Figure 1h**, **Supplementary Figure 1e**). These states were derived from three distinct molecular dynamics (MD) trajectories performed starting from the AlphaFold ORC-1 full-length model and applying diverse velocities, to maximize the exploration of the protein conformational ensemble. The considered states mainly differ from the relative orientation between the N- and C-ORC1 regions and that of the BAH domain, consistently with the prevalence of disordered and flexible residue patches between them. Both RM9 and dRM9 were docked against each of the three ORC1 states and the obtained complexes were subjected to extended molecular dynamic simulations to assess the reliability of the interaction (**Supplementary Figure 1f**). Five out of six of the ORC1-RNA complexes showed a deviation from the initial binding mode. However, the ORC1a-RM9 complex revealed a conserved protein-RNA interface and no notable structural rearrangements of either molecular entity, suggesting that RM9 efficiently targets ORC1 at the specific binding site exhibited by the ORC1a conformation. The simulation revealed the smallest perturbations from the initial state, as reflected by the lowest mean value of the root mean squared deviation (RMSD) profiles with the respect to the starting simulation state (**Supplementary Figure 1f**). Indeed, RM9 fits within the pocket embraced by the disordered region of ORC1 and the AAA+ domain, otherwise inaccessible within the 1b and 1c conformation (**Figure 1g, h**). Along with the simulation (**Tables S1-S2**), the ORC1-RM9 hydrogen indicate that RM9 mainly connects ORC1a through the ARG297, SER450, GLU575, THR593 and ARG846 residues, with occurrences greater than the 70% among the frames of the last 250 ns of trajectory (**Figure 1i**, **Supplementary Figure 1g** and **Table S2**). No significant rearrangements of the local structure of the protein were induced by the RM9 targeting, however during the simulation an approach of the BAH to the other protein domains was detected (**Supplementary Figure 1h**), suggesting that RM9 likely plays a role in favoring a specific ORC1 conformational equilibrium. We did not observe instead an effective binding mode in the ORC1a-dRM9 simulation as shown by RM9. Importantly, the simulations evidenced that the binding with DNA is facilitated by the involvement of BAH domain into the interaction and resonates with the established knowledge that the BAH domain interacts with the DNA/chromatin [27–29]. In contrast, the most efficient RM9 binding mode results from the exposure of a specific pocket in the ORC1a conformation, wherein the BAH domain is displaced away from the interacting region (**Figure 1g, h** and **Supplementary Table S1**).

Collectively, these analyses demonstrate a distinct RNA and DNA binding modality to ORC1, that support the results obtained by EMSA and RNA pull down (**Figures 1 d-f**, **Supplementary Figure 1d**).

In summary, these results pointed for a colocalization of H2A.Z and ORC1 mediated by *c-MYC ANCOR* and suggested a preferential binding of ORC1 to RNA over DNA both *in vitro* and *in silico*.

### *c-MYC ANCORs* control the replication process at the *c-MYC* locus

To assess whether alterations of the transcriptional levels of c*-MYC ANCORs* could impact the integrity of the replication process at the *c-MYC* origins we performed both CRISPR-mediated loss- and gain-of-function experiments.

Loss-of-function was achieved by repressing *c-MYC ANCORs* using CRISPR interference (CRISPRi, CRISPR-dCAS9 system)[30] in K562 cell line. Three single-guide RNAs (gRNA1-2), targeting a region located 1.4 kilobases (kb) upstream of the *c-MYC* origin, were constitutively induced, along with the scramble control, in K562 cells stably expressing dCas9-mCherry (K562 dCas9-mCherry) (**Supplementary Figure 2a,b**). The gRNAs were specifically designed to interfere with *c-MYC ANCOR* transcription without impairing with the integrity of the replication origin. The gRNA-2 led to lasting downregulation of *c-MYC* ANCOR, followed by reduction of *c-MYC* mRNA levels (**Supplementary Figure 2c**). To rule out a potential off-target effect of the CRISPRi on the *c-MYC* expression, wild type K562 were treated with FANA antisense oligonucleotides (ASOs) targeting exclusively the *c-MYC* ANCOR. The same decrease in *c-MYC* expression profile was confirmed upon reduction of the *c-MYC* ANCOR (**Supplementary Figure 2d**). Hence, the gRNA-2 was selected for all the downstream experimental analyses. In order to control the expression of the gRNAs, the selected gRNA-2 and respective scramble sequence were cloned under a Tet-inducible promoter and delivered, into K562-dCas9-mCherry, by lentiviral transduction (**Figures 2a,b**). Upon treatment with 5µM doxycycline, we followed the expression of the gRNA-2 and the scramble control over multiple time points (**Figure 2c**), and identified the peak in the gRNA levels at day three, which corresponded to the highest drop (more than two-folds) in *c-MYC ANCORs* expression (**Figure 2d**).To define the impact on DNA replication at the *c-MYC* origin, we quantified the nascent (nas) DNA abundance within the origin and found a stark decrease of nasDNA generated nearing the origin (between 2.4 to 4.5 folds), but not at the other tested regions (**Figure 2e**). Importantly, no substantial changes in cell growth as no differences in MYC protein levels between the control and the gRNA-2 expressing cells was observed upon the doxycycline treatment and after its suspension (**Supplementary Figure 2e**), suggesting that the effect on nasDNA was independent from *c-MYC* levels [31, 32].

**Figure 2.**
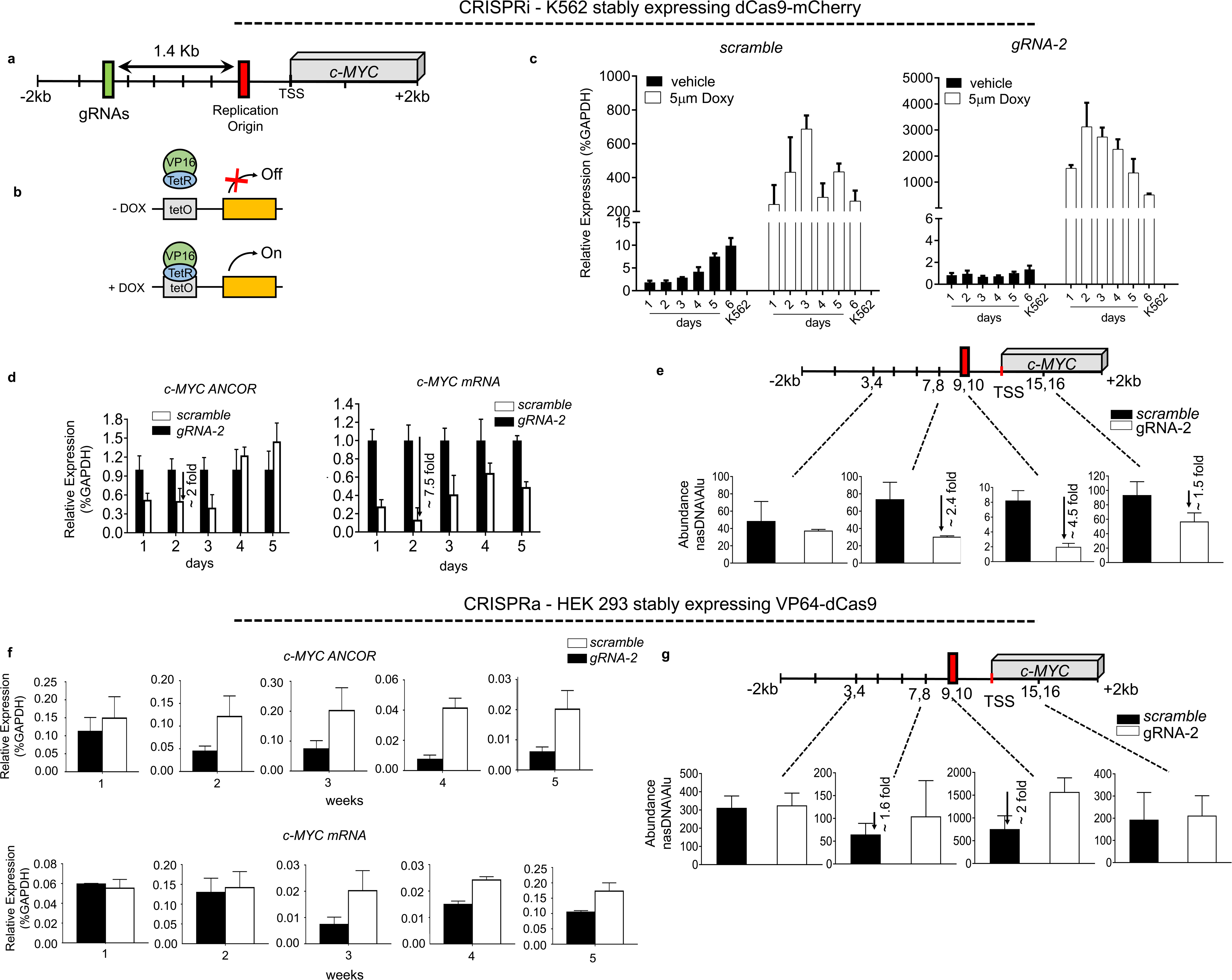
*c-MYC ANCOR* levels regulate DNA replication at the *c-MYC locus*. **a**. Outline of the *c-MYC locus* is shown, indicating the position of the guide RNA-2 (gRNA-2) sequence, located 1.4 kilobases (kb) upstream of the *c-MYC* replication origin; **b**. Schematic representation of the doxycycline-responsive promoter (TET-ON) employed to clone the gRNA-2 for the inducible CRISPR interference in K562 stably expressing dCas9-mCherry; **c**. Time-course analysis by quantitative reverse transcription (qRT)-PCR of scrambled control and gRNA-2 expression following a single 5mM doxycycline addition; **d**. qRT-PCR analysis of *c-MYC ANCOR* and *c-MYC* expression over a time course of six-days upon induction of gRNA-2 and scramble control with a single doxycycline addition; **e**. quantification of nascent (nas) DNA abundance at the *c-MYC* origin shows a significant decrease 2.4- and 4.5-folds at the regions encompassing the *c-MYC* origin:7-8 and 9-10 respectively, in response to the suppression of *c-MYC ANCOR*. Measurements were taken at day three following induction of gRNA-2 and scramble control; **f-g**. CRISPR activator (CRISPRa) in HEK 293. **f**. gRNA-2 introduced in HEK 293 constitutively expressing VP64-dCas9 results in up-regulation of *c-MYC ANCOR*, and *c-MYC* mRNA; **g**. Activation of *c-MYC ANCOR* was associated with an increase in newly synthesized DNA strands of 1.6- and 2-folds within the regions surrounding the *c-MYC* origin: 7-8 and 9-10 respectively.

Gain of function was applied using the CRISPR activator (CRISPRa, CRISPR-VP64) system [33]. To this purpose the gRNA-2 was delivered in stably expressing dCas9-VP64 HEK 293, wherein *c-MYC* is not actively replicating as in K562 [34, 35]. This led to robust up-regulation of *c-MYC ANCORs* lasting up to five weeks, paralleled by increase in *c-MYC* expression. nasDNA abundance was measured in dCas9-VP64 HEK 293 stably expressing the gRNA-2 or the scramble control. As expected, we observed enrichment of newly synthesized DNA strands at the origin of replication upon expression of *c-MYC ANCORs* as compared to the control and unlike other regions analyzed in the promoter, hence supporting the hypothesis that *c-MYC ANCORs* could promote or aid licensing and firing of *c-MYC* replication origin (**Figure 2g**). Interestingly, we also observed an increase in cell growth likely due to the increase levels of *c-MYC* protein (**Supplementary Figure 2f**).

Collectively, these results delineated the role of *c-MYC ANCORs* in the formation of *c-MYC* replication origin and underscored how alterations in their levels may perturb the integrity of the replication process at the targeted locus.

### ORC1 engagement to H2A.Z by S-Phase induced RNAs control DNA replication

The identification of *c-MYC ANCORs* and the RNA binding features retained by ORC1 and H2A.Z suggested the involvement of RNA in the formation of the replication origins and their chromatin status at the *c-MYC* locus. We, therefore sought to explore the extent of ORC1–S-Phase RNA association in other genomic loci with respect to presence of DNA replication origins. To this purpose, we assessed the transcriptional profile of S-Phase induced RNAs upon synchronization in the G1/S phase by performing nascent RNA sequencing (nasRNA-seq) at an earlier (S1) and later (S2) time point after the release on the S-phase over a course of five hours [36], (**Figure 3a**). This approach led us to the identification of 10,087 genes expressed in the earlier S1 time-point and 10171 in the later time point S2 with both including *c-MYC* within the group of genes displaying the highest expression (first quartile **Figure 3b**, **Supplementary Figure 3a**), and *FXN* among the lowest expressing genes’ group (last quartile **Figure 3b**, **Supplementary Figure 3a**). *FXN* is harbored in a mid-late replicating *locus* and its replication profile has been well characterized owing to the pathological implications in the Friedreich’s ataxia disease [9, 37, 38]. Next, we interrogated the interaction between ORC1 and the RNAs induced in the first three hours of the S-Phase to determine whether we could establish a correlation between the presence ORC1-RNA binding and the early replication origins. We uncovered 81,931 predicted loci associated with S-Ph**<u>a</u>**se RNAs a**<u>NC</u>**horing **<u>OR</u>**C1, thereby expanding on the potential global nature of transcripts showing similar features as the *c-MYC ANCORs*. Long et al. have recently proposed a function for H2A.Z in licensing DNA replication loci, and accordingly we investigated the linkage between genomic location originating *ANCORs* and enrichment of H2A.Z in the K562 cells. Over 56% loci associated with *ANCORs* overlapped with H2A.Z enriched genomic locations, accounting to 46,360 (*ANCORs*/H2A.Z) predicted early replication origins (**Figure 3c**). The *ANCOR* defined origins broadly distributed across the genome including promoters / transcription start site (TSS), gene bodies (3’ UTR, 5’ UTR, exons, and introns) and intergenic regions. Specifically, when narrowing the window for origin localization from +/−100kb to +/−0.5kb with respect to TSS, the number of predicted early DNA replication origins only decreased by two folds, from 43,188 to 19,725 with a significant proportion of these *loci* being located within the proximity of promoters and/or TSS of protein coding genes, specifically: 58% in the 2kb or 67% in the 0.5kb as compared to 28% in the 100 kb tile (**Figure 3d**, **Supplementary Figure 3b,c**). Overall, these results suggest and confirm that most DNA replication origins tend to localize in closer proximity to the promoters and/or TSS of protein coding genes [2, 39].

**Figure 3.**
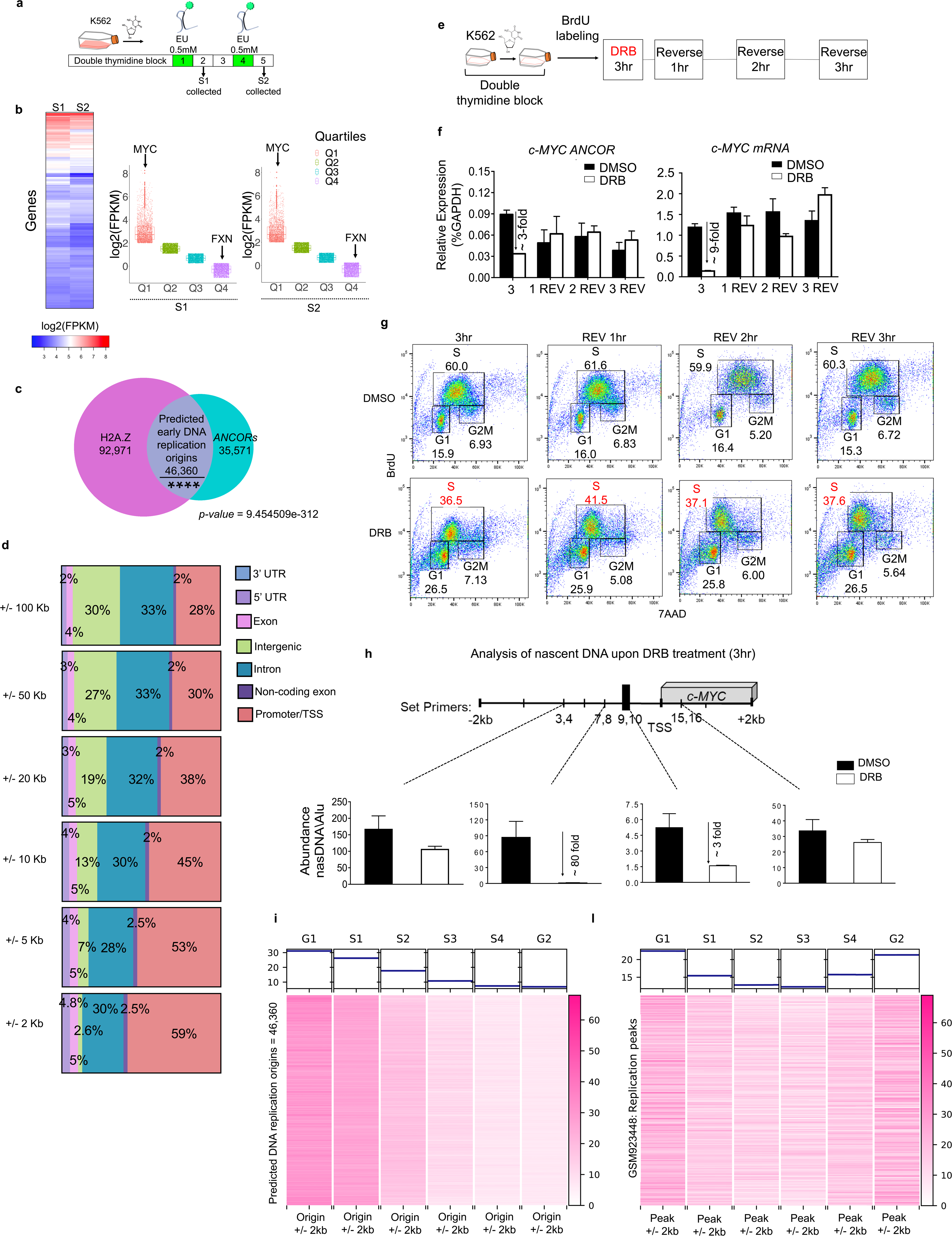
ORC1 engagement to H2A.Z by S-Phase induced RNAs mark early replication origins. **a**. Outline of the experimental setup used to identify nascent S-Phase induced RNAs. Upon synchronization in the G1/S phase by double thymidine block, the growth medium was supplemented with the EU RNA analog at the indicated time points: S1(2 hr) and S2(5 hr). EU labeled nascent RNA was collected at S1 and S2 by Click-iT conversion for RNA sequencing. **b**. Heatmap showing gene expression profiles at different stages of S-phase in K562 cells. Welch Two Sample t-test: p-value = 0.02993. Boxplot showing expression of 10,087 and 10171 genes (FPKM > 0.5) by quartiles in S1 and S2 S-phase. Expression profiling revealed *c-MYC* among the highest expressed genes (first quartile) and *FXN* among the lowest expressing genes (last quartile); **c**. Venn diagram showing intersection of S1-Phase induced RNAs anchoring ORC1 (*ANCORs*) detected by RIP-Seq and H2A.Z enriched sites by ChIP-Seq. Overlapped genomic *loci* are predicted early DNA replication origins. **d**. Annotation of the predicted early DNA replication origins located within +/−100kb, +/−50kb, +/−20kb, +/− 10kb +/−5kb and +/−2kb from the TSS. A significant proportion of the predicted early DNA replication origins lie within the proximity of promoters and/or transcription start sites (TSS) of protein coding genes; **e**. The schematic showing K562 synchronization in G1/S phase cycle followed by incorporation of BrdU into nasDNA and pharmacological inhibition of transcription using DRB (100mM). Samples were collected 3 hours after DRB treatment, and every hour upon DRB reversal over the 3 following hours; **f**. Reversible downregulation of *c-MYC ANCOR* and *c-MYC mRNA* expression by qRT-PCR upon DRB treatment and the reversal; **g**. Flowcytometry analysis showing nonreversible inhibition of the nasDNA synthesis at all selected time points; **h**. DNA abundance within the *c-MYC locus* significantly decreases upon DRB treatment only within the region encompassing the origin (7-8 and 9-10) and not the others surveyed. **i**. Heatmap showing analysis of K562 Repli-Seq signal in G1/G1b, S1, S2, S3, S4, G2 cell cycle fractions (GSE34399) centered at the ANCOR-H2A.Z predicted 46,360 early DNA replication origins in K562; **l**. Heatmap showing analysis of Repli-Seq signal centered at the replication peaks (GSE34399) in K562 cells.

To verify whether pharmacological inhibition of transcription could lead to alterations of the S-Phase in the cell cycle progression, K562 cells were synchronized by double thymidine block [35] and treated with the RNA Polymerase II (RNAPII) inhibitor DRB upon release into the S-phase. Concurrently, 3.5 µm 5-bromo-2′-deoxyruridine (**BrdU**) was added to the cells in order to monitor the nascent DNA synthesis by flowcytometry following transcriptional inhibition. We took advantage of the reversibility features of DRB and examined the effect of global transcriptional inhibition three hours upon DRB treatment and three hours after DRB reversal (treatment outline shown in **Figure 3e-g**). We monitored the transcriptional inhibition by measuring *c-MYC ANCOR* levels, three hours after DRB treatment and every hour over the three following upon DRB reversal (**Figure 3f**). A strong downregulation of *c-MYC ANCOR* and *c-MYC* mRNA (nearly 3- and 9-fold change, respectively) was noted three hours upon DRB treatment. However, these changes were transient and reversible, and the expression returned to baseline during the DRB reversal (**Figure 3f**). By contrast a non-reversible inhibition in the cell cycle progression was observed at all selected time points upon DRB reversal, as evidenced by the sharp reduction to 38% of the S-Phase cell population in DRB-treated cells as compared to the 60% S-Phase population in the DMSO-sample control (**Figure 3g**). Consistently, this pattern was associated with a parallel drop in nasDNA synthesis at the *c-MYC locus* and loss of ORC1 and H2A.Z enrichment at *c-MYC* origin upon transcriptional inhibition (**Figure 3h**, **Supplementary Figure 3d**). Similarly, DRB treatment of another cell line HL60 resulted in impairment of DNA synthesis, followed by a drastic reduction of the S-Phase cell population to nearly 30% in DRB treated cells as compared to the observed 50% in the mock control (**Supplementary Figure 3e**).

Lastly, we took the advantage of available Repli-Seq data set carried out on K562 (GSE34399) in six cell cycle fractions isolated by FACS sorting: G1/G1b, S1, S2, S3, S4, G2, to investigate the distribution of the *ANCOR-*predicted replication origins defined by the colocalization of loci associated with S-Ph**<u>a</u>**se RNAs a**<u>NC</u>**horing **<u>OR</u>**C1, and enriched in H2.AZ (**Figure 3i**). A stronger intensity of the signal was detected in the G1 and S1/S2 cell fractions when the analysis was computed with respect to the *ANCOR-*predicted replication origins, while gradually fainting in the later phases (**Figure 3i**). Considerably, when the analysis was not focused on the *ANCOR*-predicted regions but centered at the replication peaks [40], the nascent DNA signal distribution was spread across the different fractions as expected (**Figure 3l**), and consistently with the hypothesis that *ANCOR*-predicted *loci* identify early DNA replication origins.

Taken together, these data demonstrate a global interaction of S-Phase induced RNAs and ORC1, that in association with H2A.Z provide a tool for the detection of early replication origins across the genome. In parallel these findings support the hypothesis that *ANCOR*s are involved in initiation of DNA replication.

### ORC1 engagement and H2A.Z enrichment is lost upon inhibition of S-Phase induced RNAs

The changes in cell cycle progression upon DRB treatment prompted us to compare ORC1 engagement with respect to H2A.Z distribution across the 46,360 predicted early DNA replication origins (**Figure 3c**), in synchronized cells wherein pharmacological inhibition of transcription by DRB was applied in the first three hours after release from G1/S. Block of early S-Phase RNAs resulted in significant reduction of ORC1-RNAs interactions as compared to the DMSO control (*p-value* < 2.2e^−16^) and corresponded to a parallel depletion of H2A.Z (*p-value* = 4.473e^−14^) at the respective *loci* (**Figure 4a-c**). Importantly, when inspecting the *c-MYC locus* we did confirm that pharmacological inhibition of transcription using DRB resulted in depletion of ORC1-RNA interactions and of H2A.Z enrichment at the *c-MYC* origin (**Figure 4d**). This held true also for the *FXN locus* that clustered with the lowest expressing group (**Figure 3b**). Indeed, we observed the expected decrease of ORC1-RNA interactions associated with reduced enrichment of H2A.Z at the *FXN* origin upon inhibition of transcription (**Figure 4c**). Next, we stratified the K562 transcriptional profiles at the earlier (S1) and later (S2) time points after the release on the S-phase (**Figure 3a**) centering the analysis on the *ANCOR*-predicted replication origins over a tile of +/−10kb from the TSS. Consistently, we detected the higher transcriptional activity in the S1 as compared to S2 time point (*p-value* = 0.02993), corroborating the hypothesis that *ANCOR*-associated *loci* mark early replication origins (**Figure 4e**).

**Figure 4.**
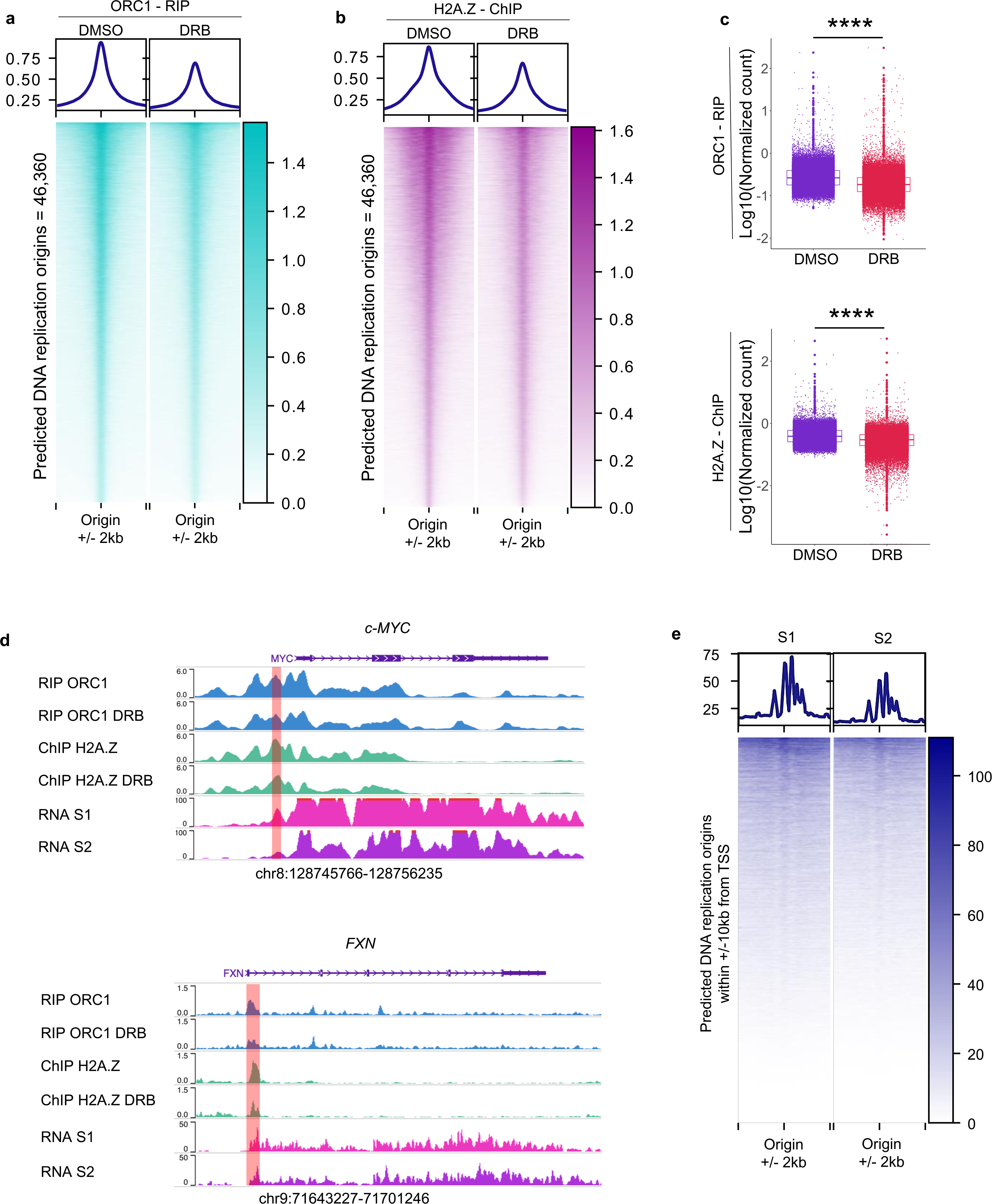
Impaired ORC1 Engagement and loss of H2A.Z Upon Inhibition of *ANCORs*. **a**. Heatmap showing decrease of ORC1 enrichment by RIP-Seq at predicted 46,360 early DNA replication origins upon DRB-treatment as compared to DMSO mock control, in K562 cells. Welch Two Sample t-test: *p-value* < 2.2e-16; **b**. Heatmap showing decrease of H2A.Z enrichment by ChIP-Seq at 46,360 early DNA replication origins in response to DRB-treatment as compared to DMSO mock control, in K562 cells. Welch Two Sample t-test: *p-value* = 4.473e-14; **c**. Box plot showing global reduction of ORC1 RIP-(upper panel) and H2A.Z ChIP- signals upon treatment with DRB in K562; **d**. Genomic snapshots of the *c- MYC* and *FXN loci* depicting the impact of pharmacological inhibition of transcription using DRB on ORC1 and H2A.Z enrichment at the respective origin sites. Transcriptional profile of the *c-MYC* and *FXN loci* for the S1 and S2 time points are presented. Higher expression of *c-MYC* and *FXN ANCORs* are observe at the S1 time point. The rose gold bar indicates the well-documented *c-MYC* origin [4, 6], and the predicted origin by the *ANCORs* association for the *FXN*; **e**. Heatmap showing the transcriptional profiles of K562 at S1 and S2 centered on *ANCOR*-predicted replication origins within +/−10kb from the transcription start site (TSS). The higher transcriptional activity in S1 supports the association of *ANCOR*-associated *loci* with early DNA replication origins.

These findings demonstrate that inhibition of transcription during the early S-Phase impairs both ORC1 engagement and H2A.Z enrichment, pointing to an RNA mediated mechanism coordinating the formation of early replication origins.

### Disruption of *ANCOR* perturbs DNA replication dynamic

In order to examine whether disruption of transcription in the S-Phase, namely reduction of *ANCORs*’ levels, would destabilize the formation of the origin and perturb the replication process, we monitored the replication fork dynamics genome-wide by leveraging DNA fiber analysis at single-molecule resolution [41, 42](**Figure 5a**). Synchronized K562 were treated with DRB for three hours upon release into the S-Phase, while sequential additions of 5-iodo-2′-deoxyuridine (IdU) and 5-chloro-2′-deoxyuridine (CldU) [42] were included into the medium to follow the progression of the replication forks during transcriptional inhibition (**Figure 5a**). Upon *ANCOR* downregulation nasDNA production was impaired and specifically, both DNA replication initiation and fork progressions were found to be decreased in K562 DRB-treated as compared to the control (**Figures 5b,c**).

**Figure 5.**
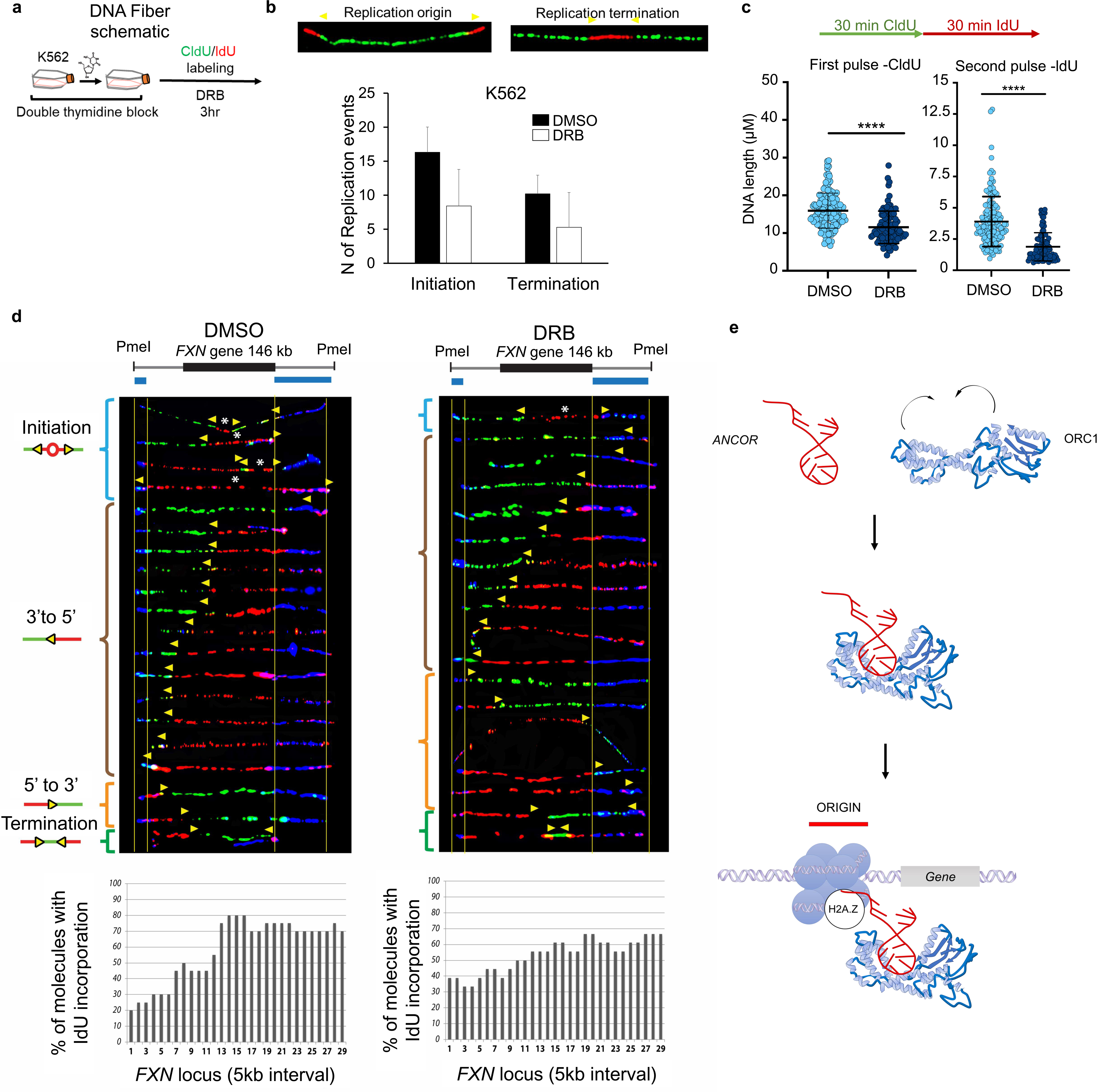
DNA Replication dynamics is perturbed by inhibition of ANCOR levels. **a**. Depicted the experimental design for DNA fiber analysis. K562 cells synchronized at G1/S phase by double thymidine block were treated with DRB for 3hrs upon release into the S phase, and sequential additions of 5-iodo-2′-deoxyuridine (IdU) and 5-chloro-2′- deoxyuridine (CldU) were used to follow the progression of replication forks during transcriptional inhibition; **b**. Quantification of DNA fibers at the “origin initiation” and “origin termination” in K562 DRB-treated cells as compared to DMSO; **c**. Quantification of DNA fibers during the two consecutive pulses with IdU and CldU, respectively for K562 DRB-treated cells compared to DMSO; **d**. Single molecule analysis of replicated DNA (SMARD) performed on *FXN locus*. Origin initiation, DNA replication fork progression (3’ to 5’ and 5’ to 3’), and origin termination were monitored upon DRB treatment and DMSO control. Percentage of molecules with IdU incorporation is indicated in the column chart below each respective condition; **e**. Proposed model for DNA origin formation. ORC1 conformational changes enable binding to RNA and formation of the complex formation (top and middle panels). The RNA engagement of ORC1 to the origin site through H2A.Z enables licensing of early DNA replication origins.

To elucidate the impact of *ANCOR* transcriptional inhibition on DNA replication progression within a specific gene *locus*, we performed SMARD (single molecule analysis of replicated DNA) on *FXN*, a *locus* amenable for this approach, unlike *c-MYC* and emerging from our analysis (**Figures 3b, 4d**) [9, 43]. Strikingly, DNA replication fork progression at the *FXN locus* was disrupted by block of transcription following DRB treatment. The abnormal replication pattern exhibited reduced initiating events, altered replication progression from 5’ to 3’ and increased terminating events when compared to the mock treatment (**Figure 5d**), thus confirming that *ANCORs* play a critical a role in maintaining the integrity of the replication process.

In conclusion, our investigation reveals that abrogation of *ANCORs* severs DNA replication program as initially suggested (**Figure 3e-g**, **Supplementary Figure 3e**), ultimately leading to a broad and deleterious perturbation of DNA replication dynamics.

## Discussion

DNA replication initiation needs to be molecularly and temporally coordinated to ensure efficient DNA synthesis and to preserve cell-specific replication dynamics.

Our investigation examines the molecular steps leading to the “*initiation*” of the DNA replication. We focused on the contribution of S-Phase induced RNAs as the masterminds coordinating the engagement of ORC1 (the first subunit of the pre-replication complex) [15] at H2A.Z enriched origins across the genome in order to license DNA replication origins in a timely and selectively fashion.

Through the identification of S-phAse-RNAs aNChoring ORC1 **– *ANCOR*s**, we delve into the role of RNA in maintaining the integrity of the DNA replication origin activity and provide evidence of the mechanism governing the initial events of DNA replication.

We demonstrate that changes in the expression of *ANCOR*’ levels result in perturbation of the replication profile and delineate a tool to regulate site-specific gene replication in human cells by means of RNA. By bringing together the first subunit of the origin recognition complex ORC1 and the histone variant H2A.Z, *ANCORs* showcase unexpected modalities to control gene replication at precise genomic sites, thereby offering an innovative approach for future translational applications. Indeed, control of gene replication “*in loco*” holds promise for precise intervention in diseases associated with dysfunctional DNA replication, wherein the lack of knowledge of how DNA replication is initiated at specific sites, has represented a major roadblock to the developmental of successful therapeutic options. Herein, we present evidence to harness RNA molecules as a tool to control *locus*-specific replication in human cells.

GC-rich elements have been identified as integral components of replication origins in human cells. These GC-rich sequences serve as binding sites for key replication factors, including the origin recognition complex (ORC) [26]. In unraveling the mechanisms governing mammalian DNA replication, we have identified a distinctive RNA sequence (RM9) embedded within the *c-MYC* origin (region 9-10) capable of binding ORC1 (**Figure 1e-g**). When compared to: 1) the unrelated and poor GC-rich RNA (*i.e.* MYC1-3) or 2) the homologous DNA (dRM9) sequences, only a weak or no interaction was detected, respectively, thus suggesting that both sequence and structural requirements were needed for the ORC1 - RNA complexation to be brought about.

By examining *in silico* a range of ORC1 conformations, mainly varying for the orientation between the N- and C-terminals, which include the bromo adjacent homology (BAH) and the AAA+ domain (ATPases associated with diverse cellular activities)/winged-helix domain (WHD), respectively, we modeled the ORC1-RNA recognition. The architecture of the whole ORC1 subunit structure remains poorly characterized owing to the presence of flexible regions, referred as disordered domains, that connect the N- and the C-terminal regions. These disordered regions within ORC1 as well as the conformational dynamics of its AAA+ domain have been extensively studied and have been considered functional to its role in inducing the replication machinery [13]. Our theoretical model demonstrates that dRM9 lies in the proximity of ORC1, when the BAH domain is part of the binding region, during the ORC1b-dRM9 and ORC1c-dRM9 simulations, while deviates from the initial position when the BAH domain does not belong to the binding site in the ORC1a state. On the contrary, the RM9 exhibits an opposite behavior, with a loss of its fold when the protein complexation involves the BAH domain and an efficient targeting when the BAH domain is displaced away from the interacting region.

Taken together these data suggest that the BAH domain is not involved in the complexation with the RNA and provide a model wherein the BAH domain needs to be structurally free to interact with the histones. In the proposed model ORC1a is the conformation that exposes a specific binding site that perfectly fits RM9 but is unable to accommodate dRM9.

Accordingly, RNA could promote a long-range effect on the ORC1 dynamic by affecting the conformational equilibrium of the protein and targeting the ORC1a conformation in a pocket formed by ORC1 disordered and AAA+ domains. This change in the conformational equilibrium would facilitate the engagement of the BAH domain for subsequent chromatin binding.

Consistently, pharmacological inhibition of transcription by DRB yielded compelling evidence demonstrating a significant global reduction of ORC1-RNA interactions and respective enrichment of H2A.Z, that was further validated at a close inspection of the *c-MYC* and *FXN loci*, using an array of approaches including fiber, SMARD analyses, in addition to sequencing. (**Figures 4–5**).

These results shed light into the interplay between transcription, chromatin dynamics, and establishment of DNA replication origins, underlining the role of RNA in the formation of active replication origins.

Our interpretation of the molecular steps involved in the DNA origin formation supports a model wherein *ANCORs* coordinate the engagement of ORC1 through the binding with H2A.Z across the genome, thereby licensing initiation early replication origins (**Figure 5e**).

Overall, our study identifies *ANCOR*s as the pacesetters for early DNA replication origins formation and alludes to a broader involvement of S-Phase induced RNAs as *initiators* of DNA replication origins.

Taken together, these data uncovered an RNA-centered mechanism to control and modulate DNA replication in eucaryotic cells at specific sites.

## Author Contributions

DGT, ADR supervised the project. ADR, SU conceived and designed the study and wrote the manuscript; SU, AKE, YZ, BQT performed experiments; LP and JL provided bioinformatic and statistical analysis of the data. TP, MD and JG performed the DNA fiber analysis and SMARD. ADR, BQT, SMM and AKE provided valuable suggestions about the project highlighting the weakness and providing possible solutions, and DGT, SMM, ADR and AKE critically reviewed the manuscript.

## Acknowledgements

This work was supported by the: DOD BMFRP Idea Development Award W81XWH-20-1-0518, the NIH/NIDDK R01DK136116, the HIRM Pilot Award 2021, NCI R00 CA188595, Italian Association for Cancer Research (AIRC) Start-Up grant #2014-15347, and Giovanni Armenise-Harvard Foundation to ADR; the award of NCI R35 CA197697, R01DK103858, W81XWH-15-1-0161, and P01HL131477-01A1 to DGT; and R50 CA211304 to AKE; National Institutes of Health under Award Number 5T32HL007917-24 to SU, the NCI K01CA222707 was awarded to BQT. SMM is supported by the National Institute of General Medical Sciences (R35GM130322) and the National Science Foundation (2153071).

We thank Dr. Luigi Vitagliano for helpful discussion, CINECA Supercomputing (ISCRA-C project: aptNAMD - HP10CDH724 and ISCRA-C: MDpapt - HP10CL0DAJ) is for computational support.

## Materials and Methods

### Cell Culture

The Chronic myeloid leukemia K562 and Acute myeloid leukemia HL-60 cell lines were procured from ATCC and cultured in RPMI-1640 media (Corning, Cat. No. 10-040-CM) supplemented with 10% FBS without antibiotics, maintained at 37 °C in a humidified atmosphere with 5% CO2. HEK293 cells obtained from ATCC were cultured in DMEM (Corning, Cat. No. 10-013-CV) with 10% FBS, devoid of antibiotics, at 37 °C in a humidified atmosphere with 5% CO2.

### RNA and DNA isolation

**Total RNA** isolation was conducted following the protocol described previously [44]. Subsequently, all RNA samples utilized in this study were subjected to DNase I treatment (10 U of Dnase I per 3 microg of total RNA) at 37°C for 1 hour, in the presence of Rnase inhibitor, to remove any contaminating DNA. To ensure complete DNA removal, the RNA samples were further purified using acidic phenol (pH 4.3). For cDNA synthesis, either Random Primers (Invitrogen) or gene-specific primers were employed in conjunction with Reverse Transcriptase (Roche Applied Science), following the manufacturer’s guidelines. The resulting cDNA was purified using the High Pure PCR Product Purification Kit (Roche Applied Science) [44].

**Nuclear RNA** was isolated using a modified method based on nuclei isolation protocols [45]. Approximately 50 million viable cells were washed with ice-cold PBS containing 5 mM vanadyl complex and 1 mM PMSF. The cells were then resuspended in ice-cold lysis buffer composed of 1x Buffer A (10 mM HEPES-NaOH pH 7.6, 25 mM KCl, 0.15 mM spermine, 0.5 mM spermidine, 1 mM EDTA, 2 mM Na butyrate), 1.25 M sucrose, 10% glycerol, 5 mg/mL BSA, 0.5% NP-40, and supplemented with various protease inhibitors, vanadyl complex, and Rnase inhibitor (RNAguard; Amersham Biosciences). The samples were incubated at 0°C for approximately 10 minutes and homogenized through multiple strokes in a Dounce homogenizer. The pelleted nuclei were further processed and centrifuged, and the resulting nuclear RNAs were extracted. All RNA samples, including total, cytoplasmic, and nuclear RNA, were treated with Dnase I to remove any contaminating DNA and then extracted with acidic phenol (pH 4.3). Polyadenylated and non-polyadenylated RNA fractions were separated using the MicroPoly(A)PuristTM purification kit (Ambion). Subsequently, cDNA synthesis was performed with Random Primers and Transcriptor Reverse Transcriptase following the manufacturer’s recommendations. The cDNA was purified using a High Pure PCR Product Purification Kit.

**Genomic DNA** isolation was performed following a previously described method [23]. Briefly, cell pellets were resuspended in lysis buffer containing 0.5% SDS, 25 mM EDTA (pH 8), 10 mM TRIS (pH 8), and 200 mM NaCl. After treatment with Rnase A (Epicentre) at 37°C for 20 minutes and Proteinase K (Roche) overnight at 65°C, high-quality genomic DNA was extracted using Phenol-Chloroform (Sigma, pH 8) and precipitated overnight with isopropanol. The genomic DNA was then resuspended in TE buffer (pH 8) and stored at 4°C.

### qRT-PCR

The Sybr Green reaction, iQ Sybr Green supermix (Biorad, Hercules, CA) was employed with the following thermal cycling parameters: an initial denaturation step at 95°C for 10 minutes, followed by 40 cycles of denaturation at 95°C for 15 seconds, annealing at 60°C for 1 minute, and extension at 72°C for 1 minute.

Regarding TaqMan analysis, Hotstart Probe One-step qRT-PCR master mix (USB) was used with the following temperature profile: an initial step at 50°C for 10 minutes, a denaturation step at 95°C for 2 minutes, followed by 40 cycles of denaturation at 95°C for 15 seconds and annealing/extension at 60°C for 60 seconds

### Primers and TaqMan probes used in real time PCR

Eukaryotic 18S rRNA Endogenous Control: ABI Cat. # 4310893E, Human GAPDH Endogenous Control: ABI Cat. # 4310884E, Human MYC mRNA: ABI Cat. # Hs00153408_m1, c-MYC ANCOR custom design (FRW-Primer:TCTGGGTGGAAGGTATCCAA, TaqMan probe:CCAACAAATGCAATGGGAGT, REW-Primer TTGGAGAGCGCGTTATGAAT)

### Primers used for strand-specific real-time RT PCR (Sybr Green)

Reverse Transcriptase primer for *c-MYC ANCOR*: 5’-AAC CGC ATC CTT GTC CTG TGA GTA-3’; PCR primers: Forward: 5’-ACA GGC AGA CAC ATC TCA GGG CTA-3’; Reverse: 5’-ATA GGG AGG AAT GAT AGA GGC ATA-3’.

### Antibodies

Rabbit polyclonal anti-ORC1 (Abcam Catalog No. ab85830); goat anti-rabbit (Invitrogen, Catalog No. 32460); rabbit polyclonal human Histone H2A.Z – ChIP Grade (Abcam Catalog No. ab4174); mouse monoclonal beta Actin (SantaCruz, sc-81178); goat anti-mouse (Invitrogen 32430); rabbit monoclonal c-MYC (Cell Signaling Technology, D84C12).

### Recombinant Proteins

Recombinant Human Origin recognition complex subunit 1(ORC1) (Cusabio Catalog N. CSB-BP621667HU(A4); mononucleosome H2A.Z (Active Motif, Cat. No. 81072)

**Nascent RNA/DNA capture** was performed using Click-iT® Nascent RNA Capture Kit (ThermoFisher) according to the manufacturer’s instructions with minor modifications. Briefly, 1. **Labeling the cells with EU/EdU**. 200 mM EU or 30 mM EdU stock solutions were added to the cells, to a final concentration 0.5 mM or 30 M, respectively. 2. **Incubation**. The cells were incubated for 1 or 2 h. 3. **RNA/DNA isolation**. The cells were harvested and the RNA/DNA were isolated and dissolved in 14 μL of H_2_O. 4. **Biotinylation of RNA/DNA by Click reaction**. Click-iT® reaction cocktail (50 μL per reaction) was prepared accordingly to manufacturer’s instructions: a mixture containing 1x Click-iT EU buffer; 2 mM CuSO_4_; 1 mM Biotin azide; 13.25 μL of the isolated RNA; 10 mM Click-iT® reaction buffer Additive 1; 12 mM Click-iT® reaction buffer additive 2 was prepared. After adding each component, the reaction cocktail was gently mixed. The addition of the Click-iT® reaction buffer Additive 1 stock initiates the click reaction between the EU-RNA/EdU-DNA and biotin azide. Subsequently, the Click-iT® reaction buffer Additive 2 is added and incubated for 30 min with gentle vortexing. 5. **RNA/DNA precipitation**. 1 μL of UltraPure™ Glycogen, 55 μL of 7 M ammonium acetate, and 750 μL of chilled 100% ethanol were added to the click reaction, incubated at –70°C for at least 30 min and after centrifugation the pellet was dissolved in 125 μL of H_2_O. 6. **Binding biotinylated RNA/DNA to Dynabeads® MyOne™ Streptavidin T1 magnetic beads (ThermoFisher)**. The RNA/DNA binding reaction mixture included: 125 μL 2xClick-iT® RNA binding buffer; 2 μL Ribonuclease Inhibitor or 2 μL of water for DNA; 125 μL of the isolated biotinylated RNA/DNA. The RNA binding reaction mixture was heated at 68–70°C for 5 min and 50 μL of bead suspension added into the heated RNA binding reaction mixture. The tube containing the RNA/DNA binding reaction was incubated at r.t. for 30 min while gently vortexing to prevent the beads from settling. The beads were immobilized using the magnet and washed 5 times with 500 μL of Click-iT® reaction wash buffer 1 and 5 times with 500 μL of Click-iT® reaction wash buffer 2. Finally, the beads were resuspended in 50 μL of Click-iT® reaction wash buffer 2 and the captured RNA immediately reverse transcribed to cDNA. The captured DNA was released into 50 μL of boiling water and used in qPCR analyses.

List of primers used to determine nasDNA abundance on *c-MYC* locus:

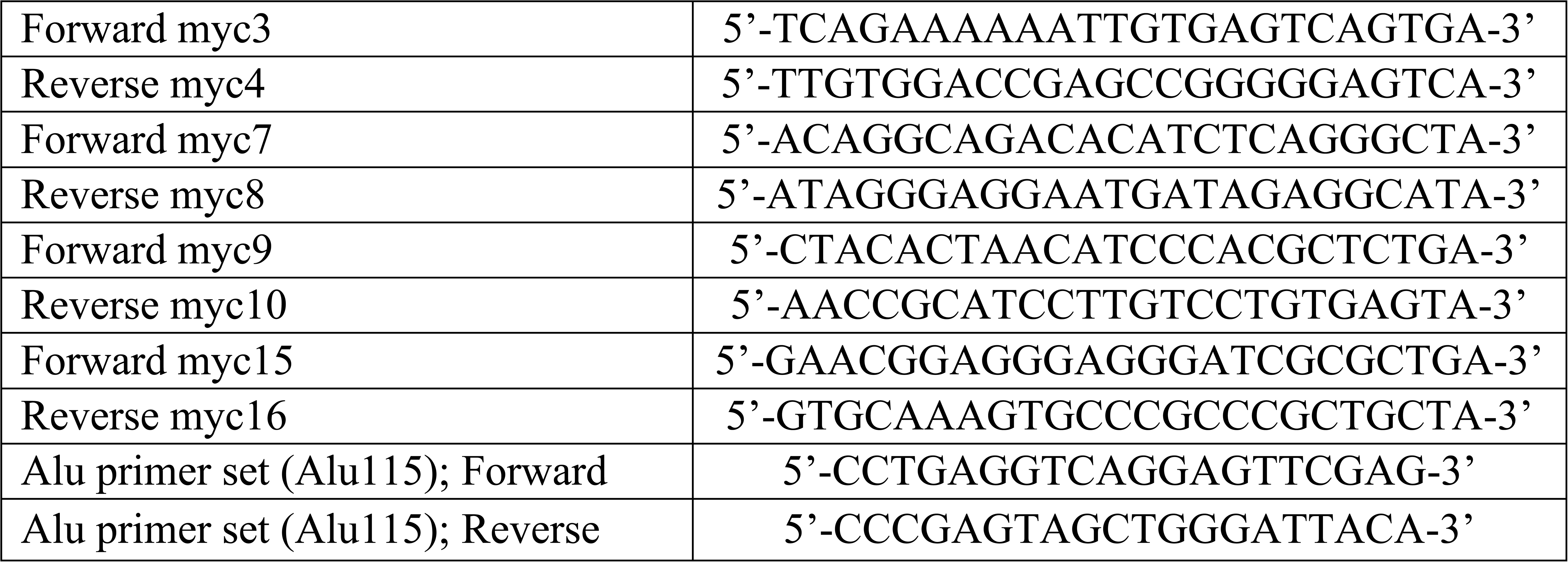

### RNA pull-down assay (RNAP)

The RNA probes were custom-designed to investigate the ORC1 binding to RNA structures transcribed within c-MYC locus including regions: c-MYC promoter, c-MYC gene and c-MYC origin. Specific set of synthetic biotinylated RNA oligonucleotides used in the RNAP is listed as follows:

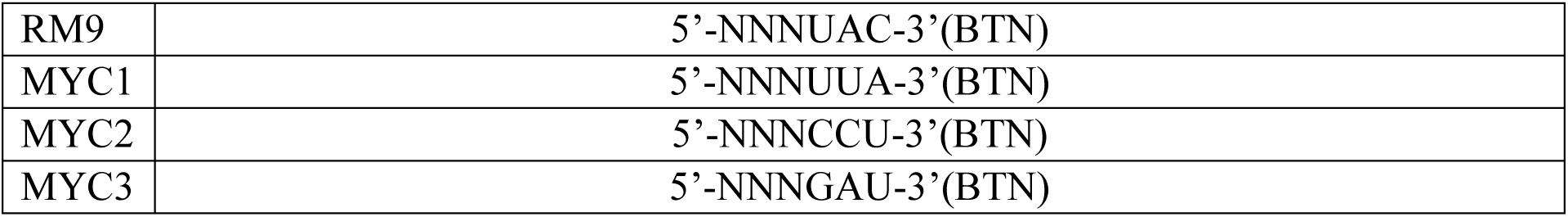

RNA-protein pull-down assay (RNAP) was conducted in triplicates following a modified protocol adapted from Pierce™ Magnetic RNA-Protein Pull-Down Kit (Thermo Scientific™, No.: 20164). Briefly, 3µg of biotinylated RNA were denatured at 90°C for 2 minutes and then allowed to fold into proper RNA structures at 25°C for 20 minutes.

3 million cells were resuspended in 1 ml of RIP Buffer containing (150 mM KCl, 25 mM Tris pH 7.4, 0.5 mM DTT, 0.5% NP40, 1 mM PMSF, and protease inhibitor (Roche Complete Protease Inhibitor Cocktail Tablets)). Cells were homogenized with a homogenizer for 20 strokes, and debris was pelleted and discarded by centrifugation at 13,000 rpm for 10 minutes. 0.1 µg of biotinylated RNA was incubated with 1 mg of cell lysate at room temperature for 1 hr. 60 µl of streptavidin agarose beads (Invitrogen) were added to the mix and incubated at RT on a shaker. The beads were then washed 10 times with RIPA buffer and boiled at 100°C in 30 μl of SDS sample buffer. Finally, the beads were removed using a magnetic field, and the samples were analyzed by western blot.

### Cell Synchronization and Drug Treatment

To induce an early S-phase block, the Double Thymidine block method was employed as previously described [35]. Briefly, K562 and HL-60 cells were grown overnight to 70-80% confluence, washed twice with 1xPBS, and then cultured in DMEM (10% FBS) supplemented with 2.5 mM Thymidine for 18 hours (first block). Subsequently, Thymidine was washed out with 1xPBS, and the cells were cultured in DMEM (10% FBS). After 8 hours, the cells were treated with Thymidine again for 18 hours (second block). Cells were then released from the block as described in reference [35]. Synchrony was assessed by flow cytometry analysis of propidium iodide-stained cells using a LSRII flow cytometer (BD Biosciences) at the Harvard Stem Cell Institute/Beth Israel Deaconess Center flow cytometry facility.

**DRB treatments,** were carried out as described [23]. Briefly, after release from double thymidine block, cells were treated with 100 μM of 5,6-Dichlorobenzimidazole 1-β–D-ribofuranoside (DRB; Sigma Aldrich).

**EU (5-ethynyl uridine, to label nasRNA) and or EdU (Ethynyl 2-deoxyuridine to label nasDNA),** were purchased from Click Chemistry Tool (CCT) and added to cell cultures at final concentration of 0.5mM (EU) or 30µM (EdU). Cells were cultured for 1hr with EU and 40min with EdU in order to reach maximum labeling of nascent RNA and DNA.

### BrdU staining and Flow cytometric Analysis of the S-Phase cycle

Minor modifications were made to the manufacture instructions provided in BD Pharmingen BrdU Flow Kit (Catalog No. 559619). Briefly, K562 cells were cultured in cell culture medium and incubated with BrdU at a final concentration of 10 µM for a pulsing period of 45 minutes. After the pulsing period, cells were resuspended in 100 µL of BD Cytofix/Cytoperm Buffer and washed with 1 mL of 1X BD Perm/Wash Buffer. The cells were then incubated with BD Cytoperm Permeabilization Buffer at all steps, keeping the samples on ice to facilitate fixation and permeabilization of the cell membranes. Following permeabilization, the cells were resuspended in 100 µL of diluted DNase, which was prepared at a concentration of 300 µg/mL in DPBS. This resulted in a dilution of 30 µg of DNase per 106 cells. The cells were incubated with DNase for 1 hour at 37°C. After the incubation, a wash step with 1 mL of 1X BD Perm/Wash Buffer was performed. To detect BrdU and total DNA content, the cells were stained with a Fluorochrome-conjugated anti-BrdU Antibody and 7-aminoactinomycin D (7-AAD) fluorescent dye. The staining was carried out for 20 minutes at room temperature The stained cells were then analyzed using a flow cytometer to determine the incorporation of BrdU and DNA content. Cell acquisition and analysis were performed on BD LSRFortessa (BD Biosciences, Franklin Lakes, NJ, USA) using BD FACSDivaTM software (BD Bioscience, Franklin Lakes, NJ, USA). Analysis was performed using Flowjo software (Flowjo LLC, Ashland, OR, USA)

### Western Blotting Analysis (WB)

**ORC1**. 0.2×10^6^ cells were lysates in SDS Sample buffer, separated on 8% SDS-PAGE gels and transferred to a nitrocellulose membrane. Immunoblots were stained overnight at 4°C with the following primary antibodies: anti-ORC1 (1:1000 TBST/5% milk; Abcam, ab85830). Immunoblots were then incubated in secondary goat anti-rabbit (1:1000 TBST/5% milk; Invitrogen, 32460) 1h at room temperature. Immuno-reactive proteins were detected using the Pierce^®^ ECL system (Thermo Scientific #32106).

**H2A.Z**. Whole-cell lysates from approximately 0.2×10^6^ cells per sample were separated on 13% SDS-PAGE gels and transferred to a nitrocellulose membrane. The membrane was blocked with TBST/5% BSA for 4 hours. Immunoblots were stained overnight at 4°C with the following primary antibodies: H2A.Z (1:2,000; Abcam ab4174), Secondary goat anti-rabbit (Invitrogen, 32460), diluted 1:4000 and incubated for 1h at room temperature with TBST/5% BSA. Immuno-reactive proteins were detected using the Pierce^®^ ECL system (Thermo Scientific #32106).

**c-MYC.** Cells (0.2×106) were lysed using SDS Sample buffer and the lysates were then subjected to separation on 8% SDS-PAGE gels, followed by transfer to a nitrocellulose membrane. The membrane was incubated overnight at 4°C with the primary antibody mAb c-MYC (dilution 1:1000 in TBST/5% milk; Cell Signaling Technology, D84C12). Subsequently, the membrane was incubated with secondary goat anti-rabbit antibodies (dilution 1:1000 in TBST/5% milk; Invitrogen, 32460) for 1 hour at room temperature. Immuno-reactive proteins were visualized using the Pierce® ECL system (Thermo Scientific #32106).

### Electrophoretic gel mobility shift assays (EMSAs)

DNA and RNA oligonucleotides were end-labeled with [γ-^32^P] ATP (Perkin Elmer) and T4 polynucleotide kinase (New England Biolabs). Reactions were incubated at 37 °C for 1 h and then passed through G-25 spin columns (GE Healthcare) according to the manufacturer’s instructions to remove unincorporated radioactivity. Labeled samples were gel-purified on 10% polyacrylamide gels. Binding reactions were carried out in 10 µl volumes in the following buffer: 5 mM Tris pH 7.4, 5 mM MgCl_2_, 1 mM DTT, 3% v/v glycerol, 100 mM NaCl. 5 nM of full length purified ORC1 (Cusabio Catalog N. CSB-BP621667HU(A4) and H2A.Z (Active Motif, Cat. No. 81072) proteins were incubated with 1.1 nM of ^32^P-labeled single-stranded (ss) RNAs. All reactions were assembled on ice and then incubated at room temperature for 30 min in Binding Buffer (HEPES-NaOH 25mM, Magnesium Acetate 10mM, Sodium Glutamate 100mM, EDTA 0,1 mM, DTT 1mM, ATP 3mM, RNase 20U, Glycerol 5%) [26]. Samples were loaded onto 6% native polyacrylamide gels (0.5xTBE) at 4 °C and ran for 3h at 140 V. All gels were fixed (20% Methanol, 10% Acetic acid), dried and exposed to X-ray film. RNA oligonucleotides are listed in **Supplementary Table B**.

### Chromatin immunoprecipitation (ChIP) and Nascent Chromatin immunoprecipitation (nasChIP)

ChIP was performed as follows. Cells were crosslinked with 1% formaldehyde for 10 min at room temperature. Pellets of 1×10^6^ cells were used for immunoprecipitation as previously described [22, 23] and lysed for 10 min on ice and chromatin fragmented using a Branson 250 digital sonicator. Each ChIP was performed with 4ug of antibody, incubated overnight at 4°C. A 50/50 slurry of protein A and protein G Dynabeads was used to capture enriched chromatin, which was then washed before reverse-crosslinking and proteinase K digestion at 65C. Immunoprecipitated DNA was analyzed by ChIP-PCR analysis and ChIP-seq. The following antibodies were used for ChIP: H2A.Z (Abcam ab4174, lot GR3176820-1), and IgG (Abcam ab171870). Fold enrichment was calculated using the formula 2^(-ΔΔCt(ChIP/non-immune^ ^serum))^.

Primer sets used for ChIP are listed in **Supplementary Table A**.

### RNA immunoprecipitation (RIP)

The RIP protocol was conducted following the established procedure [23] with slight modifications. In summary, 1.6×10^6^ K562 cells were crosslinked with 1% freshly made formaldehyde solution: 50 mM HEPES-KOH; 100 mM NaCl; 1 mM EDTA; 0.5 mM EGTA; 11% formaldehyde) for 10 min at room temperature. 2. Crosslinking was stopped by adding 1/10^th^ volume of 2.66 M Glycine, kept for 5 min at room temperature and 10 min on ice. 3. Cell pellets were washed twice with ice-cold PBS (freshly supplemented with 1 mM PMSF). 4. Cell pellets were resuspended in cell lysis buffer (volume = 4 mL): 1x Buffer (10 mM Tris pH 7.4; 10 mM NaCl; 0.5% NP-40), freshly supplemented with protease inhibitors (protease inhibitors cocktail: Roche Applied Science, Cat. No. 1836153, 1 tablet/375 μL H_2_O; add as ×100), 0.1 mM PMSF, and 0.2 mM vanadyl complex (NEB). 5. Cells were incubated at 0°C for 10-15 minutes and homogenized in a Dounce (10 strokes pestle A and 40 strokes pestle B). 6. Nuclei were recovered by centrifugation at 2,000 rpm for 10 min at 4°C. 7. Nuclei were resuspended in 3 ml of 1x Resuspension Buffer (50 mM HEPES-NaOH, pH 7.4; 10 mM MgCl_2_) supplemented with 0.1 mM PMSF and 0.2 mM vanadyl complex. 8. DNase I treatment (250 U/ml) was performed for 30 min at 37°C, and EDTA (final concentration 20 mM) added to stop the reaction. 9. Resuspended Nuclei were sonicated once for 20s (1 pulse every 3 seconds) at 30% amplitude (Branson Digital Sonifer, Danbury, CT).

**Immunoprecipitation** was performed as follows: 1. Before preclearing, the sample was adjusted to 1% Triton X-100; 0.1% sodium deoxycholate; 0.01% SDS; 140 mM NaCl; Protease inhibitors; 0.2 mM vanadyl complex; 0.1 mM PMSF. 2. Preclearing step: ∼ 50 μL magnetic beads (Protein A or G Magnetic Beads; #S1425S or #S1430S NEB) were added to the sample and incubation was carried out for 1 h on a rocking platform at 4°C. 3. Beads were removed in the magnetic field. 4. The sample was then divided into aliquots according to different conditions and antibodies of interest: (1) H2A.Z (Abcam, ab4174), ORC1 (Abcam, ab85830), preimmune serum: IgG (ab171870); (v) no antibody, no serum (input). 5. 5 μg antibody or preimmune serum was added to the respective aliquot and incubation performed on a rocking platform overnight at 4°C. Input was stored at −20 °C after addition of SDS to 2% final concentration. Day II. 6. 200 μL of Protein A coated super-paramagnetic beads (enough to bind 8 μg IgG) were added to the samples and incubated on a rocking platform for 1 h at 4°C. 7. Six washes of beads in the magnetic field were made with immunoprecipitation buffer (150 mM NaCl; 10 mM Tris-HCl, pH 7.4; 1 mM EDTA; 1 mM EGTA pH 8.0; 1% Triton X-100; 0.5% NP-40 freshly supplemented with 0.2 mM vanadyl complex and 0.2 mM PMSF) in a magnetic field. 8. Proteinase K treatment to release DNA/RNA into solution and to reverse the crosslinking was performed in 200 μL of: 100 mM Tris-HCl, pH 7.4; 0.5% SDS for the immunoprecipitated samples and in parallel for the input using 500 μg/mL of Proteinase K at 56°C overnight. 9. Day III. Beads were removed in the magnetic field. 10. Immunoprecipitated RNAs were further purified by extraction with Phenol (pH 4.3) after addition of NaCl (0.2 M final concentration). 11. Ethanol precipitation (in the presence of glycogen); 3 h at −20°C. 12. The pellet was dissolved in 180 μL H_2_O, heated at 72 °C for 2 min, and immediately chilled on ice.13. Samples were treated with DNase I (20 units) in the presence of RNase inhibitor at 300 U/ml in x1 buffer # 2 (NEB) at 37°C for 30 min. 14. Phenol (pH 4.3) extraction and EtOH precipitation were repeated. 15. The RNA pellet was dissolved in 50 μL H_2_O and used in qRT-PCR (Primer sets used for RIP-PCR are listed in **Supplementary Table A**).

### ASO/FANA-ASO AntiSense Oligonucleotides-mediated downregulation

ASOs targeting c-MYC *ANCOR* and no-targeting scrambled control were purchased from FANA AUM biotech, proprietary sequence. They were added to cell cultures at 10-20 µM concentration. Downregulation was monitored, and cell samples were collected at 48, 72, and 96 hours after ASO treatment for subsequent qPCR analysis.

### Cell Culture and Transfection

The K562 and HEK 293 cell lines were maintained in appropriate culture media supplemented with 10% fetal bovine serum (FBS) and 1% penicillin-streptomycin before lentiviral transduction. The guide RNA (gRNA) targeting the c-MYC ANCOR in a region located 1.4 kilobases (kb) upstream of the c-MYC origin were designed using the online tool (https://portals.broadinstitute.org/gpp/public/analysis-tools/sgrna-design), and an initial guide screening was conducted.

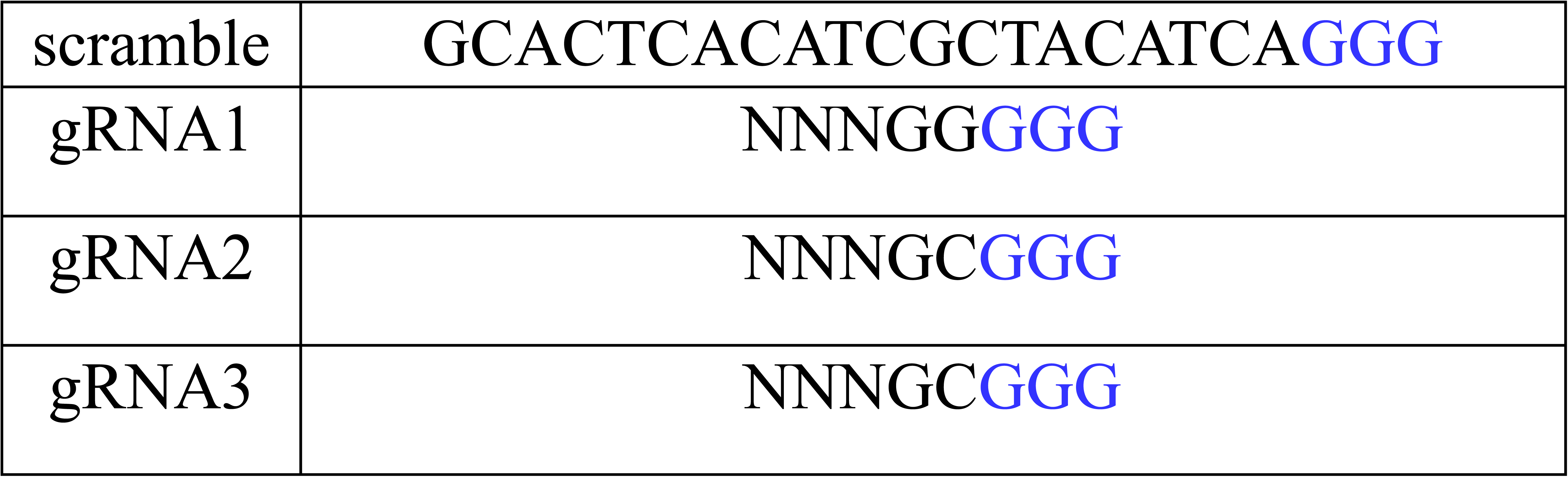

### Down-regulation of c-MYC ANCOR

#### dCAS9-mediated stable and inducible downregulation

To implement CRISPR interference (CRISPRi), the lentiviral vector pdCAS9-humanized (code 44246) with an engineered mCherry fluorescent signal was acquired from Addgene and transduced into K562 cells, following the protocol outlined in Larson MH et al (36713455). Four lentiviral vectors: Cellecta-pRSGT16-sgRNA containing guide RNAs and no-targeting control guide (gRNA-1, gRNA-2, gRNA-3, ntRNA) were purchased from Collecta, packaged into second-generation lentiviral particles, and transduced into K562 cells stably expressing dCas9 using Lipofectamine 3000 transfection reagent (Thermo Fisher Scientific) according to the manufacturer’s instructions. In the Tet-inducible system, all guide RNAs were cloned under the control of a Tet-inducible promoter. In parallel, a non-targeting RNA control was also transfected. To induce the expression of gRNAs, cells were treated with a single addition of 5µM doxycycline. For assessing guide RNA-2 expression in cells, custom primers and TaqMan probe were designed, the list is provided below:

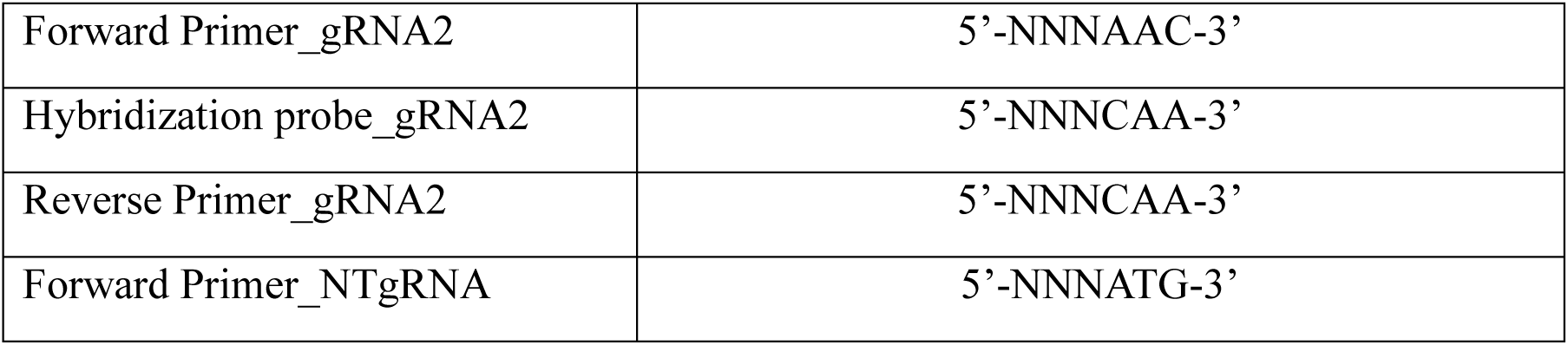

### Up-regulation of c-MYC ANCOR

#### dCAS9_VP64-mediated upregulation

*A* lentiviral vector (pLenti-U6-sgRNA-PGK-Neo) encoding the designed gRNA was synthesized by Applied Biological Materials Inc. pMD2.G, pCMV delta R8.2, and the lentivector were transfected into 10 million 293T using TransIT-LT1 reagent (Mirus). Supernatants were harvested at 48 and 72 hrs after transfection. Virus was concentrated 100 times by Lenti-X Concentrator (Takara) after filtering through a 0.45 syringe filter, and resuspended in DMEM and stored at –80^0^C. dCas9-VP64 (Addgene, plasmid #61425) lentiviral transduction was performed in the presence of hexadimethrine bromide (final concentration 8 g/ml) in HEK 293. Blasticidin (10 g/ml) was added to the cultures 2 days after infection. Stably expressing dCas9-VP64 cells were subsequently transduced with *c-MYC ANCOR* gRNA2 or scrambled control. G418 (500 g/ml) was added to the cultures 2 days after infection. Resistant clones were selected and screened to asses *DNA replication and c-MYC ANCOR levels*.

#### DNA fiber Analysis

K562 cells were cultured and maintained in appropriate growth media until reaching a cell number of 15 million. Synchronization in the S phase of the cell cycle was achieved using the Double Thymidine block method, following established procedures [35]. Additionally, DRB treatments were employed to induce an early S-phase block in transcription, as described in the literature[23]. DNA fiber assay was performed as previously described [46]. Briefly, cells were sequentially pulse labeled with 25 µM CldU followed by 250 µM IdU for 30 minutes at 37℃. Untreated cells were used as controls. Cells were washed with ice-cold PBS, trypsinized, and resuspended at 2.5×10^6^ cells/mL concentration in cold PBS. 2µl of cell suspension was gently placed on glass slide and after incubation of 2-3 min, lysis buffer (200 mM Tris-HCl; pH 7.5, 50 mM EDTA and 0.5% SDS) was added, followed by 2 min of incubation at room temperature. The slides were tilted at an angle (15-45°) to allow the fibers to spread along the slide. After drying, the slides were fixed in methanol-acetic acid (3:1) for 15 minutes at −20℃, denatured in 2.5 M HCl for 30 minutes, and blocked in 5% BSA in 1X PBS for 20 min (blocking buffer). Primary anti-BrdU antibodies specific for CldU (1:15) and IdU (1:15) were applied for 1 hour at room temperature followed by PBS wash. Slides were then stained with secondary antibodies; anti–rat Alexa Fluor 488 (1:15) and anti–mouse Alexa Fluor 568 (1:15) for 1 hour at room temperature. Slides were mounted with ProLong Antifade Mountant (Thermo Fisher Scientific #P10144,) and imaged on Carl Zeiss Axio Imager M2 microscope. Labeled replication structures were identified and the track length was measured using ImageJ. The replication events-initiation and termination were also recorded and quantified. We used student t-test with two-tailed distribution for p-value calculation.

#### Single molecule analysis of replicated DNA (SMARD) Analysis

When the cells reached 70-80% confluence, they were synchronized in S phase using the Double Thymidine block method [35] and DRB treatments were employed to induce block of early S-phase transcriptional activities as described in the literature [23].

The cells were then incubated at 37°C for 4 h in the presence of 25 μM 5-iodo-2′-deoxyuridine (IdU; Sigma-Aldrich, St. Louis, MO). After washing cells with PBS, hESCs medium with 25 μM 5-chloro-2′-deoxyuridine (CldU; Sigma-Aldrich, St. Louis, MO) was added to the cultures, and the cells were incubated for an additional 4 h. The cells were harvested with Accutase for hESC or Trypsin for mammalian epithelial cells. Following centrifugation, the cells were resuspended at 3 × 10^7^ cells per ml in PBS. 1% InCert agarose (Lonza Rockland, Inc, FMC, lot 221291) in PBS was added to an equal volume of cells at 42°C. The cell suspension was pipetted into a chilled plastic mold with 0.5- by 0.2-cm wells with a depth of 0.9 cm for preparing DNA gel plugs. The gel plugs were allowed to solidify on ice for 30 min. Cells were lysed in buffer containing 1% *n*-lauroylsarcosine (Sigma Aldrich), 0.5 M EDTA, and 20 mg/ml proteinase K. The gel plugs remained at 50°C for 64 h and were treated with 20 mg/ml proteinase K, every 24 h. Gel plugs were then rinsed several times with Tris-EDTA (TE) and once with phenylmethanesulfonyl fluoride. The plugs were washed with 10 mM MgCl_2_ and 10 mM Tris-HCl (pH 8.0).

The genomic DNA in the gel plugs was digested with 40 units of *PmeI* (at 37°C overnight. The digested gel plugs were rinsed with TE and cast into a 0.7% SeaPlaque GTG agarose gel (Lonza Rockland, Inc.). A gel lambda ladder PFG marker and yeast chromosome PFG marker (both from New England BioLabs, Inc.) were cast next to the gel plugs. PCR was performed to determine the appropriate positions of DNA on the pulsed-field electrophoresis gel and the gel piece was cut and melted at 72°C for 20 min. β-Agarase enzyme (1 unit per 50 μl of agarose suspension) was carefully added to digest the agarose and incubated at 45°C for a minimum of 2 h. The resulting DNA solutions were stretched on 3-aminopropyltriethoxysilane coated glass slides. The DNA was pipetted along one side of a coverslip that had been placed on top of a silane-treated glass slide and allowed to enter by capillary action. The DNA was denatured with sodium hydroxide in ethanol and then fixed with glutaraldehyde.

The slides were hybridized overnight with a biotinylated probe (the blue bars diagrammed on the maps indicate the positions of the probes used). The following day, the slides were rinsed in 2× SSC buffer with 1% SDS and washed in 40% formamide solution containing 2 × SSC at 45°C for 5 min and rinsed in 2 × SSC-0.1% IGEPAL CA-630. Following several detergent rinses (4 times in 4× SSC-0.1% IGEPAL CA-630), the slides were blocked with 1% BSA for at least 20 min and treated with Avidin Alexa Fluor 350 (Invitrogen Molecular Probes) for 20 min. The slides were rinsed with PBS containing 0.03% IGEPAL CA-630, treated with biotinylated anti-avidin D (Vector Laboratories) for 20 min, and rinsed again. The slides were then treated with Avidin Alexa Fluor 350 for 20 min and rinsed again, as in the previous step. The slides were incubated with the IdU antibody, a mouse anti-bromodeoxyuridine (Becton Dickinson Immunocytometry Systems), the antibody specific for CldU, a monoclonal rat anti-bromodeoxyuridine (anti-BrdU) (Accurate Chemical and Scientific Corporation) and biotinylated anti-avidin D for 1 h. This was followed by incubation with Avidin Alexa Fluor 350 and secondary antibodies, Alexa Fluor 568 goat anti-mouse IgG (H+L) and Alexa Fluor 488 goat anti-rat IgG (H+L) (both from Invitrogen Molecular Probes), for 1 h. After a final PBS/ IGEPAL CA-630 rinse, the coverslips were mounted with ProLong Gold Antifade Mountant. A fluorescent microscope (Axioscop 2 M2 with Plan Apochromat 63×/1.4 NA oil differential interference contrast objective; Carl Zeiss) with a camera (CoolSNAP HQ; Photometrics) was used to detect the nucleoside incorporation into the DNA molecules. Images were processed with Photoshop CS5 software. Image processing involves alignment of the DNA molecules according to the FISH probes, adjusting of the contrast, and removing unspecific background signal. The molecules presented are the complete dataset.

For *FXN* gene locus, biotinylated probes were prepared from Fosmids G248P83100G6 and G248P85473G3 were selected using the public gene database (http://genome.ucsc.edu/).

To analyze the replication fork progression, the % molecules with IdU incorporation moving across *FXN* locus were quantified. The segment was divided into 5kb fragments and replication fork per segment was enumerated. The percentage of molecules with IdU incroporation was calculated for the 5kb interval.

### Modeling and Molecular Dynamics simulations

The RM9 and dRM9 models of were built by using the MC-sym/MC-fold and the 3dnus procedures [47]. The coordinates of the Orc1 protein as deposited in the alphaFold protein structure database were downloaded and used as starting model of the entire protein, because no experimental solved structures are available in the pdb data bank contain the full-length protein coordinates. The created theoretical models of the interacting systems were individually subjected to molecular dynamic simulations to refine the structures and to explore the conformational dynamics, following the protocols already validated. In details, 100ns of simulations time were used to minimize RM9 and dRM9 systems, assessing the validity of their initial conformations. For what concern the Orc1 alphaFold predicted structure we applied three diverse velocities simulations in order to refine the model and better explore the Orc-1 conformational ensembles, which contains a high level of flexibility. Indeed, the high prevalence of disordered regions particularly between the N-(residues 1-479) and the C-(residues 480-861) segments suggest that the protein can explore various equally probable states and no experimental data are available to define the relative orientations between these domains. We thus extracted from each of the three trajectories a representative ORC1 conformation (namely orc-1a, orc-1b and orc-1c) and used each as target to dock the RM9 and dRM9 sequences. The pdb 7JPS (PubMed id: 32808929) structure were used as template to accommodate the RNA sequence. In details, orc-1a orc-1b and orc-1c states were structurally overlapped to the Orc-1 subunit of 7jJPS (chain A) pdb and the RNA sequence models positioned to replace the region occupied by the DNA molecules of the same pdb, that was used as template for the docking.

### Illumina-based RNA sequencing (RNA-Seq)

Strand-specific transcriptome libraries were constructed on BGI Genomics platform (DNBSEQ). Pipeline of the experiment: 1) mRNA and non-coding RNAs are enriched by removing rRNA; 2)by using the fragmentation buffer, the mRNA and non-coding RNAs are fragmented into short fragments (about 200∼700 bp); 3) the first-strand cDNA is synthesized by random hexamer-primer using the fragments as templates; 4) buffer, dNTPs, RNase H and DNA polymerase I are added to synthesize the second strand cDNA; 5) the double strand cDNA is purified with QiaQuick PCR extraction kit and then used for end-polishing; 6) sequencing adapters are ligated to the fragments and then the second strand is degraded using UNG (Uracil-N-Glyosylase); 7) the fragments are purified by Agarose gel electrophoresis and enriched by PCR amplification. Libraries were paired-end sequenced on Hi-Seq-4000 Illumina. Raw reads were analyzed using a pipeline for RNA-seq analysis - Visualization Pipeline for RNA-seq analysis (VIPER) based on workflow management system Snakemake (https://github.com/hanfeisun/viper-rnaseq). The read alignment to hg19 reference genome was performed using STAR aligner (2.7.0f) with default parameters. Gene expression (FPKM values) was quantitated with Cufflinks (v2.2.1). Bedtools (v2.27.1) genomecov and bedGraphToBigWig v 4 were used to generate bigwig files.

### RIP-sequencing

RNAs were processed for sequencing as described by Di Ruscio *et al*. with minor adaptations. RNA samples were depleted of ribosomal RNA with Ribo-ZeroTM Magnetic Gold Kit (cat. # MRZG126 Epicentre). Libraries were constructed on BGI Genomics platform (DNBseq, RIPseq). Pipeline of the experiment: 1) RNA was enriched; 2) the first-strand cDNA was synthesized by random hexamer-primer using the fragments as templates 3) following by second strand synthesis and ER/A-tailing; 4) ligation with an overhanging T at the 3’ end of the bubble adapter; 5) PCR amplification of the ligated products with specific primers; 6) single-strand separation by thermal denaturation and circularization of the single-stranded DNA through a bridge primer to obtain single-stranded circular DNA library; 7) sequencing (amplified products DNA Nanoballs (DNBs) were used for paired-end sequencing on Hi-Seq-4000 Illumina. Raw data with adapter sequences or low-quality sequences was filtered. This step was completed by SOAPnuke software developed by BGI. SOAPnuke software filter parameters:-n 0.001 -l 20 -q 0.4 -A 0.25 --cutAdaptor -Q 2 - G --minLen 100. Raw reads were analyzed using a pipeline CHromatin enrIchment ProcesSor Pipeline (CHIPS) based on workflow management system Snakemake (https://github.com/liulab-dfci/CHIPS) where the read alignment to hg19 reference genome was performed using bwa mem aligner (0.7.15-r1140) with default parameters; MACS2 algorithm (2.2.7.1) was used to call peaks; and MACS2 (2.2.7.1) and bedGraphToBigWig (v 4) were used to generate bigwig files. Read counts for downstream analyses were quantified using the bioliquidator/bamliquidator tool from Docker image (https://hub.docker.com/r/bioliquidator/bamliquidator). Genomic regions were annotated using Hypergeometric Optimization of Motif EnRichment tool (HOMER v3.12).

### ChIP Sequencing

ChIP-libraries were constructed on BGI Genomics platform (DNBseq, CHIPseq). ChIP library construction pipeline: 1) end repair and adapter ligation (filling the ends of IP/Input DNA followed by 5’ phosphorylation. A 3’ sticky-end with an overhanging A is formed, which could ligate with an overhanging T at the 3’ end of the bubble adapter); 2) PCR amplification (PCR amplification of the ligated products with specific primers); 3) single-strand separation and circularization (separation of PCR products into single-strands by thermal denaturation. Then circularization of the single-stranded DNA through a bridge primer to obtain single-stranded circular DNA library); 4) sequencing (amplified products DNA Nanoballs (DNBs) were used for paired-end sequencing on Hi-Seq-4000 Illumina. Raw data with adapter sequences or low-quality sequences was filtered. We first went through a series of data processing to remove contamination and obtain valid data. This step was completed by SOAPnuke software developed by BGI. SOAPnuke software filter parameters:-n 0.01 -l 20 -q 0.4 --cutAdaptor -Q 2 -G --polyX 50 --minLen 100. Raw reads were analyzed using a pipeline CHromatin enrIchment ProcesSor Pipeline (CHIPS) based on workflow management system Snakemake (https://github.com/liulab-dfci/CHIPS) where the read alignment to hg19 reference genome was performed using bwa mem aligner (0.7.15-r1140) with default parameters; MACS2 algorithm (2.2.7.1) was used to call peaks; and MACS2 (2.2.7.1) and bedGraphToBigWig (v 4) were used to generate bigwig files. Read counts for downstream analyses were quantified using the bioliquidator/bamliquidator tool from Docker image (https://hub.docker.com/r/bioliquidator/bamliquidator). Genomic regions were annotated using Hypergeometric Optimization of Motif EnRichment tool (HOMER v3.12).

### Graphics

Genomic Heatmaps were generated using computeMatrix and plotHeatmap scripts from deeptools (v3.5.1). All Boxplots were generated using the “ggplot2” R package (https://ggplot2.tidyverse.org). ScatterDensity plots were generated using the smoothScatter R function (https://www.rdocumentation.org/packages/graphics/versions/3.6.2).

Genome browser pictures were generated using the “WashU Epigenome Browser” (https://github.com/lidaof/eg-react/)

### Statistical Analysis

All the statistical analyses were performed using the R suite (https://www.r-project.org/). The statistical comparison of the distributions corresponding to ChIP-Seq, RIP-Seq and RNA-Seq cumulative read count signals were performed using the Mann Whitney Wilcoxon signed rank test, with a confidence interval of (℘ < 0.05). For the statistical evaluation of HAT assay, we applied two-way ANOVA with multiple comparisons. GraphPad Prism Software was used. Values of p≤ 0.05 were considered statistically significant.

## Data Availability

Sequencing Data are available on the gene omnibus database under the accession ID number: GSE241684.

---

**Supplementary Figure 1.**
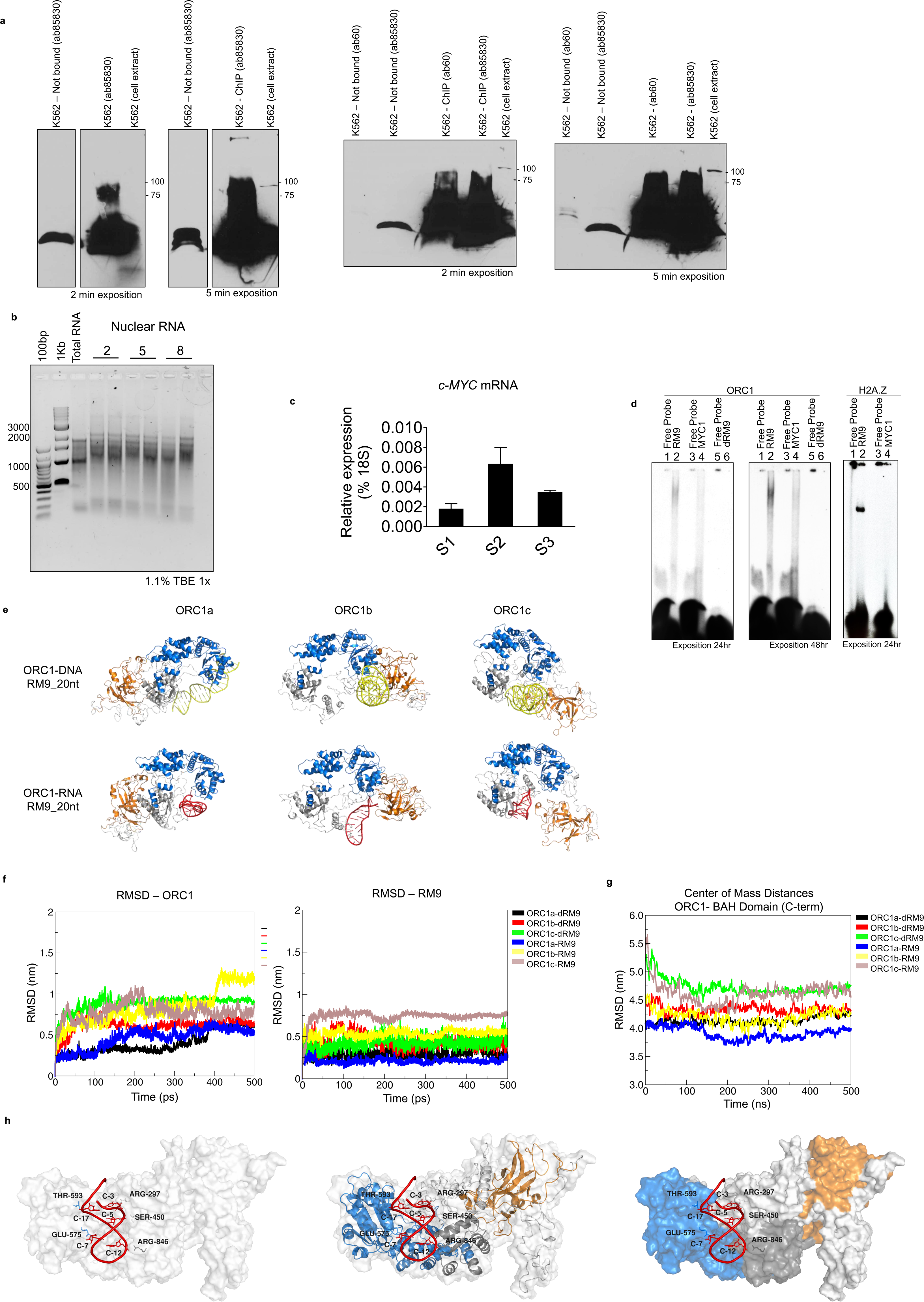
Characterization of ORC1-S-Phase induced RNA interaction. **a**. Validation by western blot analyses of two independent ORC1 antibodies using either K562 whole cell extract and ORC1 immunoprecipitates; **b** Electrophoresis analysis shows the integrity of total RNA and nuclear RNA fractions run on 1.1% Agarose gel in 1x Tris-Borate-EDTA (TBE) buffer; **c**. Expression of *c-MYC* mRNA at different time points of the S-phase of the cell cycle (S1, S2 and S3 respectively 2, 5 and 8 hours) performed on nuclear RNA; **d**. RNA and DNA Electrophoretic Mobility Shift Assays (EMSA) show binding of ORC1 to RM9 (RNA), but not MYC1 or the homologous DNA sequence dRM9 (left and middle panels indicate different exposure time). RM9 interacts with ORC1 unlike MYC1 (right panel); **e**. Frontal view of three apo ORC1 states (ORC1a, ORC1b, and ORC1c) showing distinct conformations, mainly differing in the relative orientation between the N- and C- ORC1 regions and the BAH domain. orange: ORC1-BAH domain (residues 1-200), marine ORC1 disordered region and AAA+ domain (residue 201-782), gray: orc1-WHD domain (residues 783-861); red and yellow RM9 and dRM9, respectively; **f**. Root Mean Squared Deviation (RMSD) profiles of the ORC1-RM9 complexes during extended molecular dynamics (MD) simulations. The ORC1a-RM9 complex exhibits the smallest perturbations from the initial state. **Left panel**: The RSMD profiles computed using only the C-alpha (C-a) atoms of each ORC1 state residues along 500 ns of six complex MD simulations with the respect to the starting run state. Orc1a-dRM9, black (mean: 0.6 nm, SD: 0.15), orc1b-dRM9, red (mean: 0.6 nm, SD 0.09), orc1c-dRM9, green (mean: 0.9 nm, SD: 0.09), orc1a-RM9, blue (mean: 0.5 nm, SD: 0.13), orc1b-RM9, yellow (mean: 0.8 nm, SD: 0.21) and orc1c-RM9, brown (mean 0.9 nm, SD 0.11). **Right panel**: The RSMD profiles computed using only the C1’ atom of either RNA or DNA sequence along 500 ns of six complex MD simulations with the respect to the starting state. Orc1a-dRM9, black (mean: 0.3 nm, SD: 0.05), orc1b-dRM9, red (mean: 0.4 nm, SD 0.09), orc1c-dRM9, green (mean: 0.4 nm, SD: 0.07), orc1a-RM9, blue (mean: 0.2 nm, SD: 0.03), orc1b-RM9, yellow (mean: 0.5 nm, SD: 0.04) and orc1c-RM9, brown (mean 0.7 nm, SD 0.07);; **g**. Distances between the center of mass of ORC1-BAH domain (residues 1-200) and the C- terminal region (200-861) along the entire simulations. Orc1a-dRM9: black, orc1b-dRM9: red, orc1c- dRM9, green orc1a-RM9: blue, orc1b-RM9: yellow and orc1c-RM9: brown; **h**. Hydrogen bond interactions between RM9 and ORC1a residues, indicating the key residues involved in the interaction. ARG297, SER450, GLU575, THR593, and ARG846 predominantly mediate the connection between ORC1a and RM9 during the simulation.

**Supplementary Figure 2.**
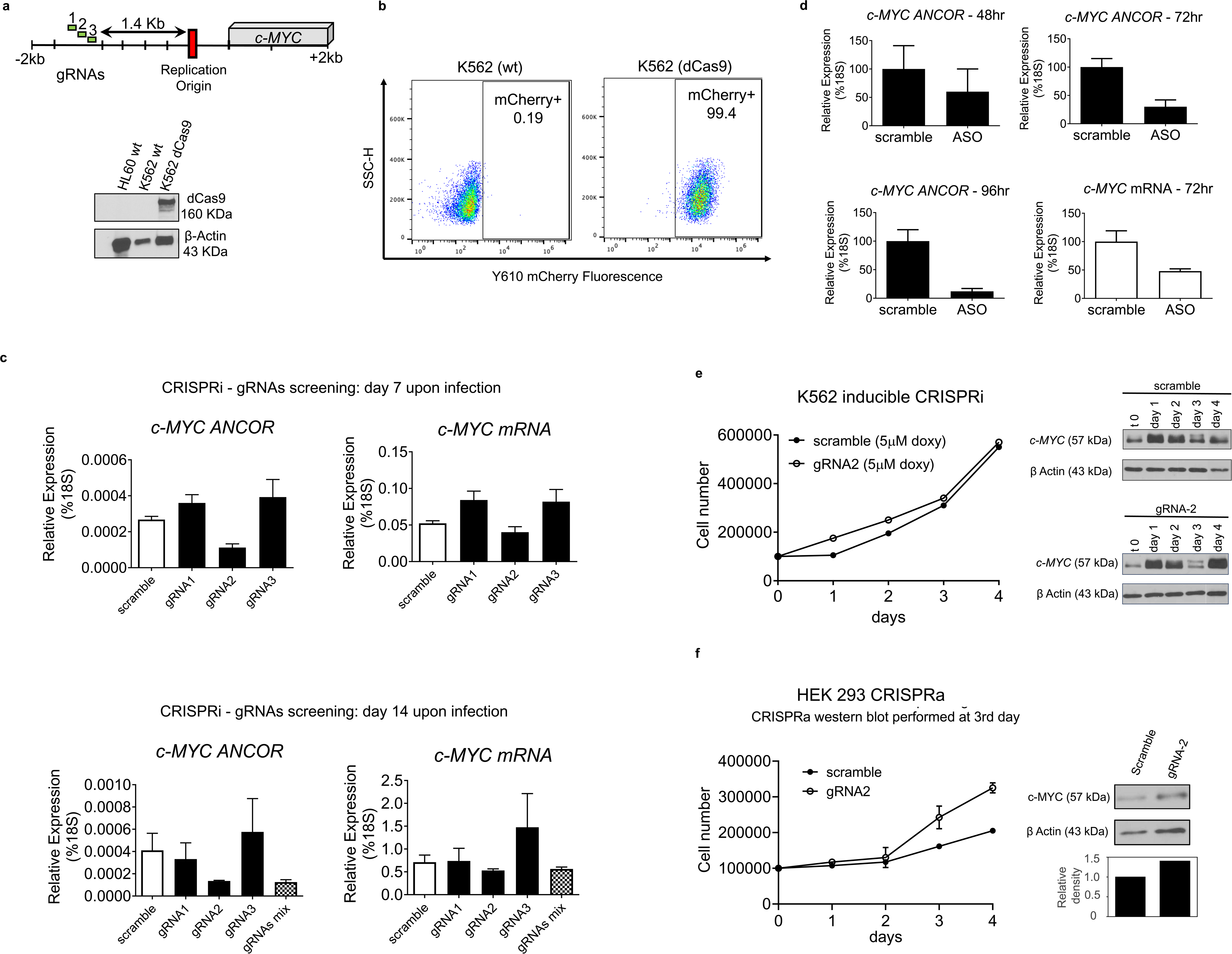
*c-MYC ANCOR* loss and gain of function by CRISPRi and CRISPRa. **a**. Schematic of *c-MYC* locus showing gRNAs targets and expression of dCas9 protein levels in K562 cells assessed by western blot analysis; **b**. Flow cytometry analysis showing percentage of K562- expressing dCa9-mCherry (99.4%); **c**. qRT-PCR screening the effect of non-inducible gRNAs on *c-MYC ANCOR* and *c-MYC* mRNA levels, 7 and 14 days upon viral transduction in K562-dCa9-mCherry.; **d**. ASOs targeting *c-MYC ANCOR* and a non-targeting control were delivered into K562 at a concentration between 15 µM. Samples collected at 48, 72 and 96 hours were used for qRT-PCR. Downregulation of *c-MYC ANCOR* was observed starting from 48 hours, lasting until 96 and was associated with significant reduction of *c-MYC* levels **e**. Growth curve of K562 dCas9-mCherry, expressing the inducible gRNA-2 or the control sequence. No changes in cell number were observed upon induction of gRNAs by doxycycline in the first four days (left panel). Immunoblot analyses of c-MYC levels across a four days’ time-course were comparable between the gRNA-2 and scramble control transduced cells (right panel); **f**. Growth curve of HEK 293 stable cells expressing dCas9- VP64 with gRNA-2 or the scramble control over a four days’ time-course. Higher cell count was measured at day3 in the gRNA-2 transduced cells as compared to the control (left panel) and parallel increase of c-MYC protein was detected by western blot at day 3.

**Supplementary Figure 3.**
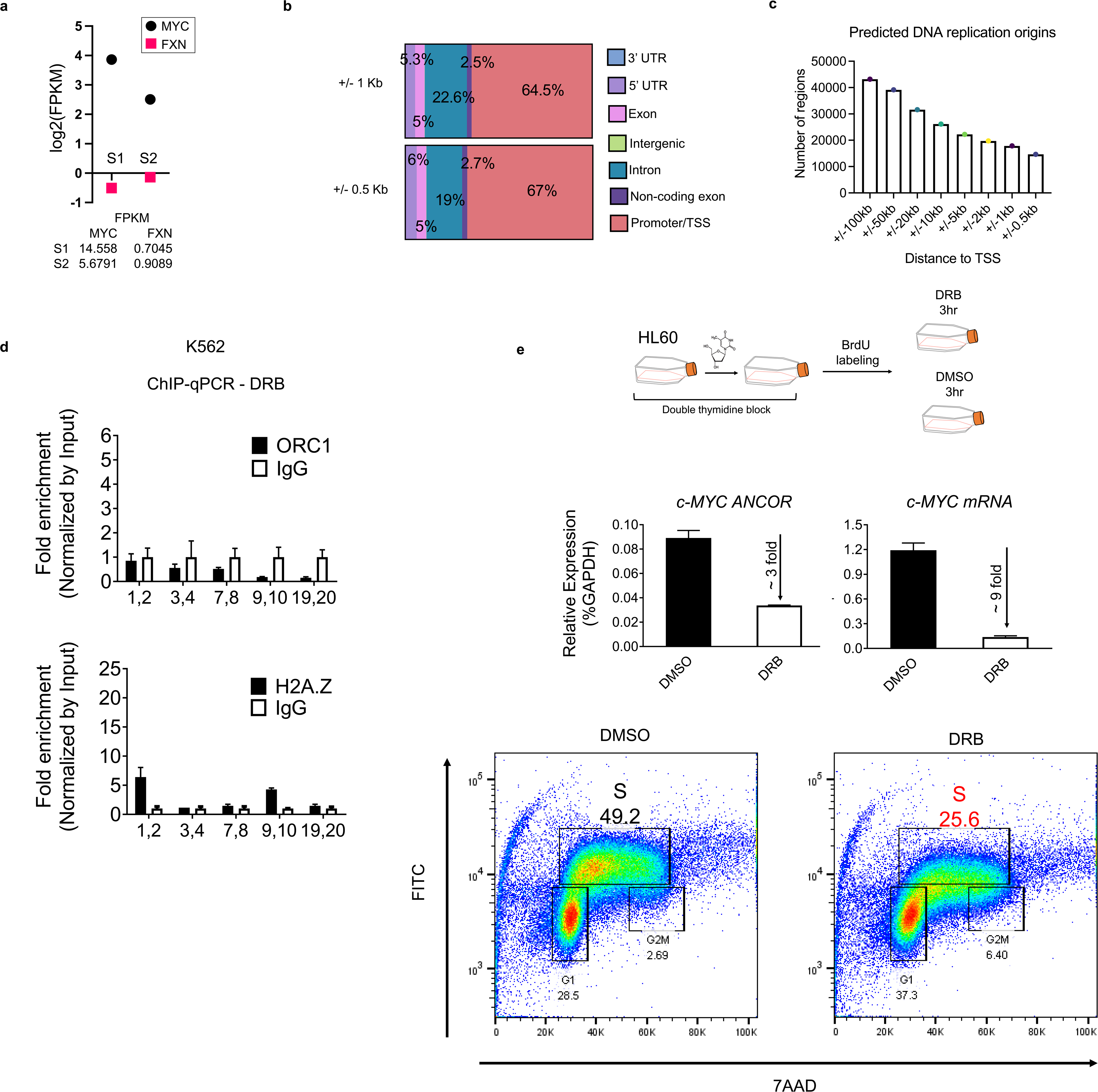
I S-Phase induced RNAs control DNA replication through ORC1 and H2A.Z. **a**. RNA-seq analysis depicting the gene expression levels (FPKM) of *c-MYC* and *FXN* at earlier (S1) and later (S2) stages of the S phase; **b**. Annotation of predicted early replication origins across the genome demonstrated that a substantial portion lies in close proximity to promoters and/or transcription start sites (TSS) of protein-coding genes. The annotation highlights the predicted DNA replication origins positioned within +/−1kb and +/−0.5kb from the TSS; **c**. Histogram graph illustrating the distribution of predicted early DNA replication origins, represented by the number of examined regions plotted against the distance from the transcription start site (TSS); **d**. Chromatin immunoprecipitation and qPCR analyses (ChIP-qPCR) following DRB (100µM) treatment show decreased enrichment of ORC1 and H2A.Z at the *c-MYC* origin (region 9-10). The qPCR analysis was performed with different primer sets, as indicated on the x-axis; **e**. (Upper panel) Schematic representation of HL60 cells synchronization by double thymidine block into G1/S phase, followed by BrdU (3.5 µm) addition in the growth medium to label nascent DNA, and either pharmacological inhibition of transcription by DRB (100µM) or mock treatment with DMSO. Samples were collected 3-hours upon DRB treatment and control. (Middle panel) qRT-PCR analysis showing downregulation of *c-MYC ANCOR* and *c-MYC* mRNA expression upon DRB treatment. (Bottom panel) Flowcytometry analysis depicting inhibition of the nasDNA synthesis with a drastic reduction of the S-Phase cell population to nearly 30% in DRB treated cells as compared to the observed 50% in the DMSO control.

**Supplementary Table A.**
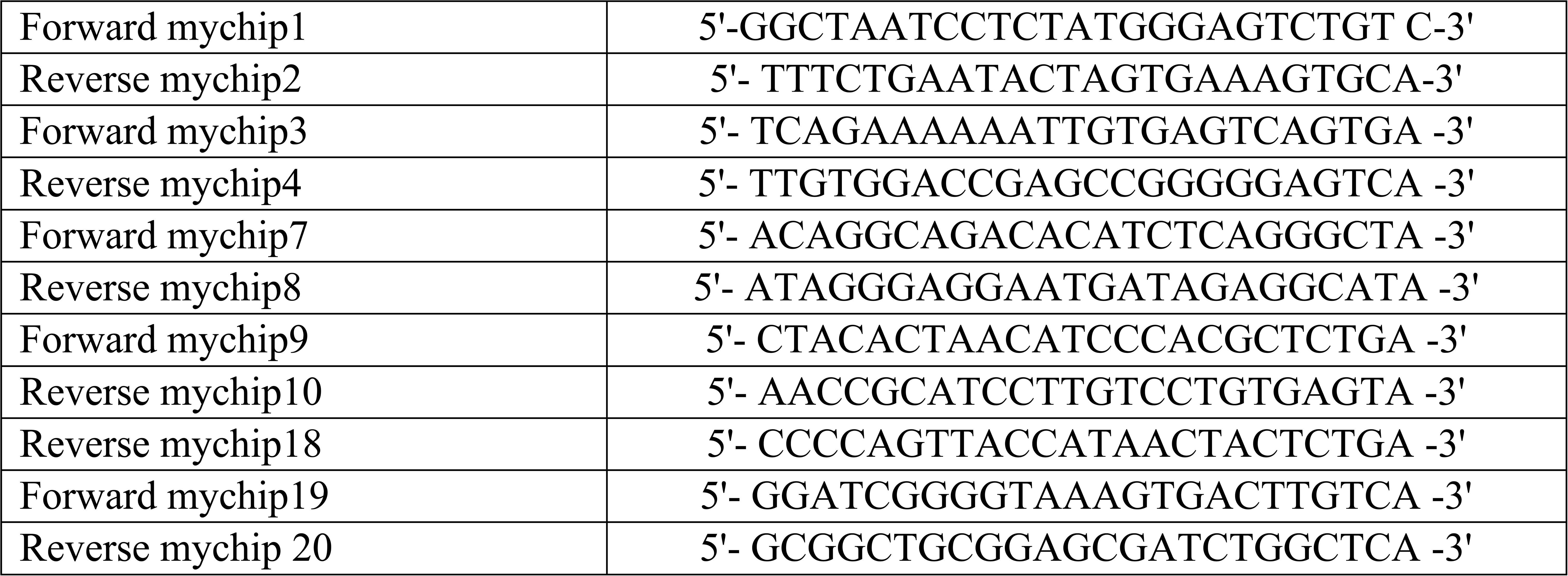
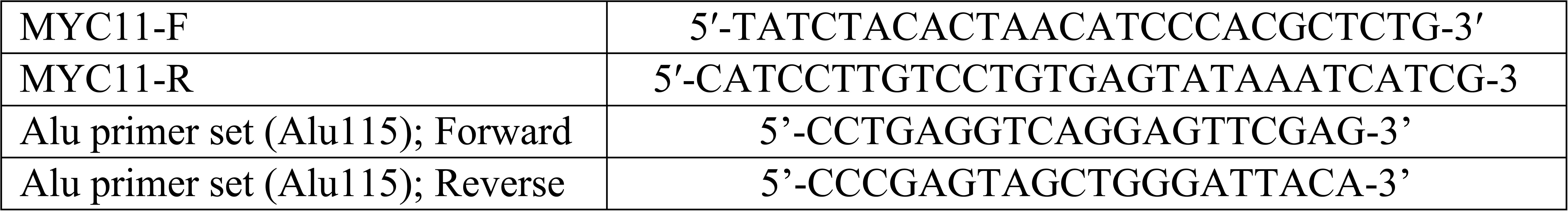
Primer-Sequences used for Chromatin and RNA Immunoprecipitation qPCR (*c-MYC* locus)

**Supplementary Table B.**
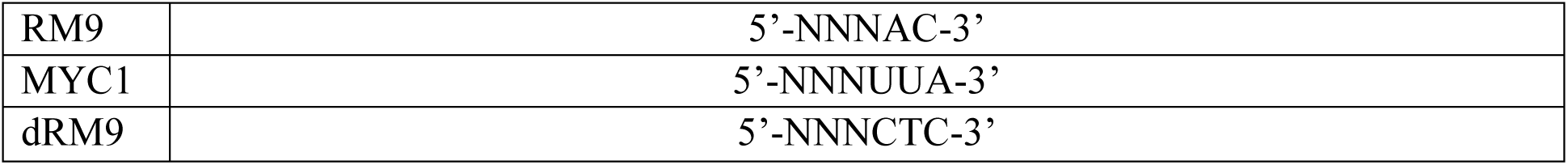
RNA/DNA Oligonucleotides used in EMSA assay:

